# Oncogenes have the most distinct codon biases in the genome and codon signatures that oppose tumor suppressor genes

**DOI:** 10.64898/2026.08.21.746283

**Authors:** Chetna Mathur, Evan T. Davis, Dylan Ehrbar, Humphrey C. Omeoga, Lauren Endres, Shane R. Byrne, Ulrike Begley, Peter C. Dedon, Thomas J. Begley

**Author notes:** Correspondence: Thomas J. Begley. The authors declare they have no actual or potential competing financial interests.

## Abstract

Oncogenes and tumor-suppressor genes play opposing roles in cancer biology to promote and restrict growth, respectively. Codon usage patterns interface with tRNA modifications to control translation, leading to gene-specific codon signatures with regulatory potential. As such, codon-biased translational regulation has been identified as a driver of proliferation and drug resistance in multiple cancers. We used advanced codon analytics methods to characterize and compare codon usage bias in oncogenes and tumor suppressor genes (TSGs) from humans and mice at group and gene-specific levels. We demonstrate that human oncogenes exhibit a distinct and opposing codon usage pattern to TSGs. This phenomenon is also present in mice but with less distinct oncogene bias relative to humans. Further comparison to 447 gene ontology groups demonstrated that human oncogenes have the most distinct codon usage patterns in the genome, while also highlighting that codon bias can separate functionally related genes and pathways from other biological processes. Using gene-specific codon analytics, we determined that human oncogenes have two types of extreme codon bias: a large group (N = 43) over-using G/C ending (GC3) codons and a smaller group (N = 12) over-using A/U (AU3) ending codons. While GC3 bias has been linked to increased translation in general, the AU3 finding suggests that genetic, environmental, or stress-related signals could drive the translation of this small group of oncogenes. The less extreme bias observed in mouse oncogenes and tumor suppressors likely underscores species-specific differences in oncogenic translation programs. Together, our findings highlight codon usage bias as a potential determinant of oncogene expression, provide a framework for ontology-based codon analysis, and uncover on species-specific differences in oncogene translation and codon usage biases.

## 1. Introduction

Cancer is a complex disease characterized by uncontrolled growth that can drive the progressive transformation of cells into malignancy (Hayden et al., 2008). Cancers can arise due to genetic mutations or epigenetic changes that disrupt normal cellular processes and gene regulation associated with appropriate growth, survival and death. Genetic alterations, including gene amplifications and deletions that result in copy number variations (CNVs), can dramatically affect cellular hemostasis and promote cancer onset and progression, especially when occurring in oncogenes and tumor suppressor genes (Hanahan et al., 2011). Active protein synthesis is required to fuel the rapid proliferation of cancer cells, and translational controls are crucial in modulating gene expression signature associated with signatures in cancer (Endres et al., 2019; Robichaud et al., 2019; Kovalski et al., 2022; Añazco-Guenkova et al., 2024). Several oncogenes are intricately linked to the regulation of translation, influencing protein synthesis and contributing to tumorigenesis. Overexpression of eIF4E has been observed in cancers such as acute myeloid leukemia (AML), breast cancer, and prostate cancer, leading to enhanced translation of oncogenic mRNAs with complex 5’ UTRs (Carroll et al., 2013). MYC enhances ribosome biogenesis and translation by upregulating components of the translational machinery, including rRNA, ribosomal proteins, and translation initiation factors. Overexpression of MYC is implicated in various cancers, such as Burkitt’s lymphoma, breast cancer, and lung cancer (Dhanasekaran et al., 2022). Translational dysfunction is central to cancer and more than 100 other human diseases (Torres et al., 2014; Jonkhout et al., 2017; Rapino et al., 2017; Barbieri et al., 2020; Dedon et al., 2022) highlighting the importance of translational control to human health (Hershey et al., 2012).

The process of translation uses transfer RNA (tRNA) and several translation factors in the ribosome to decode messenger RNA (mRNA). Maintaining the integrity and regulation of this system is essential for protein synthesis. Alongside tRNA aminoacylation, numerous base modifications are essential for regulating tRNA activity (El Yacoubi et al., 2012; Suzuki et al., 2021), and are catalyzed by specific writer enzymes (Boccaletto et al., 2022). Writers play a key role in RNA structure, function, and codon-decoding specificity. They also influence translational fidelity, gene expression, splicing, RNA degradation, and translational regulation (Wang et al., 2014; et al., 2015; Xiao et al., 2016; Meyer et al., 2017; Shi et al., 2019). Post-transcriptional RNA modifications are essential for regulating various cellular processes, including differentiation, activation, migration, mRNA development, and polarization (Roundtree et al., 2017; Barbieri et al., 2020; Cui et al., 2022). The location and type of chemical modifications on tRNAs impact function, including tRNA stability and codon-anticodon interactions that dictate translational efficiency (Väre et al., 2017; Boccaletto et al., 2022). Notably, modifications at the tRNA wobble position are crucial for optimizing anticodon-codon interactions during translation, and disruptions are linked to mitochondrial and neurodevelopmental disorders, with altered wobble uridine (U) and corresponding writer levels linked to cancer proliferation and chemotherapeutic resistance (Ashraf et al., 1999; et al., 2000; Agris et al., 2004; et al., 2007; Rapino et al., 2018; Orellana et al., 2021).

Degeneracy of the genetic code allows multiple synonymous codons to serve a regulatory function in cells, which means that more than one codon can code for the same amino acid. Codon-biased translation involves the enhanced translation of certain codons and corresponding mRNAs due to the increased availability or affinity of tRNAs with complementary anticodons. tRNA can enhance the production of codon-biased mRNAs that encode proteins that respond to cellular metabolic conditions and environmental stressors, as well as promote tumor growth by enhancing codon-anticodon interactions (Chan et al., 2012; Gu et al., 2014; Zhang et al., 2022). For instance, the writer tRNA methyltransferase 1 (METTL1), together with its cofactor WDR4, methylates the guanine (G) at position 46 in the variable loop of specific tRNAs, converting it to 7-methylguanosine (m^7^G). METTL1 is increased in multiple tumors and drives the production of codon-biased oncoproteomes (Dai et al., 2021; Orellana et al., 2021; Han et al., 2022). Across diverse human cancers, *METTL* family members which are either amplified or upregulated with consistently poorer clinical outcomes (Campeanu, 2021). Additionally, the ELP3/CTU1/2 complex, which is essential for wobble uridine (U) modifications), has been shown to drive melanomas with the BRAF^V600E^ mutation, leading to resistance to anti-BRAF therapy due to codon-biased translation of metabolic proteins (Rapino et al., 2018). Both METTL1 and ELP have been associated with enhanced translation of mRNAs possessing specific codon usage patterns (Chan et al., 2012; Roundtree et al., 2017; Dedon et al., 2022). METTL1-mediated m^7^G tRNA methylation preferentially enhances translation of mRNAs enriched in codons decoded by m^7^G -modified tRNAs, including GAA (Glu), CUC (Leu), and AAC (Asn) (Lin et al., 2018; Orellana et al., 2021), while ELP3/CTU1/2-driven wobble U modifications promote efficient translation of AA-ending codons (e.g., GAA, AAA, CAA) decoded by U_34_-modified tRNAs, thereby reinforcing codon-biased expression programs in tumors (Ladang et al., 2015; Delaunay et al., 2016; Rapino et al., 2018). Codon-biased translation is driven by the increased availability and affinity of tRNAs with chemically modified anticodon bases, which enhances translation of the specific cognate codons and corresponding mRNAs. Initially observed in the regulation of stress response in yeast corresponding mRNAs are called modification-tunable transcripts (MoTTs) (Chan et al., 2012; Dedon et al., 2014) and have been identified in bacteria, viruses, mice, and humans (Chionh et al., 2016; Dedon et al., 2022; Jungfleisch et al., 2022; Mitchener et al., 2023).

Codon usage bias has been recognized as an important determinant of gene regulation, and here we report that oncogenes and tumor suppressors (TSGs) have opposing codon patterns in humans and mice, with more pronounced differences in humans than that of mice. We report that human oncogenes exhibit the most extreme codon bias of 447 other defined gene ontologies (GOs) and TSGs. Oncogenes further segregate based on codon usage biases into, a small group that overuses A/U-ending codons (AU3), and a larger group which favors G/C-ending codons (GC3). The GC3 group, includes FGF4, SKI, and CCND1 genes. We also compared murine oncogenes to TSGs and 462 other mouse GOs and determined that mouse oncogenes also display extreme codon usage biases. However, the magnitude of this bias is more pronounced in human oncogenes. Our findings reveal codon usage bias as a distinctive feature of oncogenes, provide a blueprint for strategies that targeting oncogene translation in specific cancers, and highlight a key difference in oncogene codon bias between humans and mice.

## 2. Material and methods

### 2.1 Gene datasets

Human oncogenes and TSGs were compiled from the Cancer Genes website (http://cbio.mskcc.org/CancerGenes/Select.action) (Walker et al., 2012). Genes whose status as either oncogenes or TSGs remained uncertain or still questionable were omitted from the final list. For humans, we assembled a list of 73 TSGs and 66 oncogenes. We also assessed our results using an independently curated gene set published in 2024 (Trexler et al., 2024), which consisted of 54 oncogenes and 71 TSGs (Supplementary data). Mouse TSGs (55) and oncogenes (45) were obtained from Uniprot (http://www.uniprot.org/keywords) (Consortium et al., 2025). We also included 447 human and 462 mouse gene ontology-based groupings (all ontologies with between 50 to 150 genes) found in the AmiGO2 browser (http://amigo.geneontology.org) (Ashburner et al., 2000; Carbon et al., 2009; Aleksander et al., 2023). Gene Ontology (GO) enrichment analysis of a list of human and mouse genes was performed to identify significantly over-represented biological processes, molecular functions, and cellular components. This type of enrichment analysis by AmiGO2 is usually performed using Gene Ontology annotations, which are accessed by AmiGO2 and is called over-representation analysis (ORA). We also compared the codon bias usage of TSGs and oncogenes between humans and mice. The lists of oncogenes and TSGs for both humans and mice are shown in **Table 1 and 2.**

### 2.2 Codon Analytics

Oncogenes and TSGs were analyzed using both isoacceptor codon frequency (ICF) and total codon frequency (TCF) metrics. ICF describes the occurrence of synonymous codons relative to each amino acid and was determined by dividing the number of occurrences of each codon for a specific amino acid by total number of codons for the amino acid within a specific gene **(Formula 1)**.

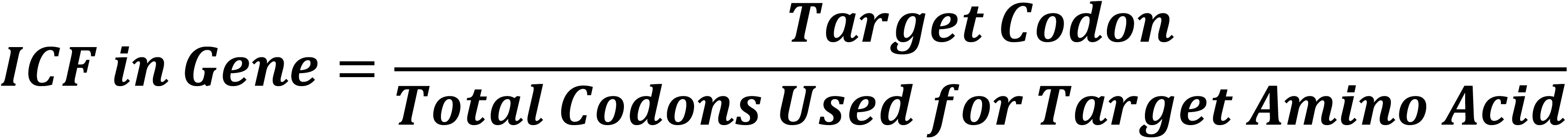

TCF was determined by dividing the number of times a specific codon was found in a gene by the total number of codons in that same gene **(Formula 2)**.

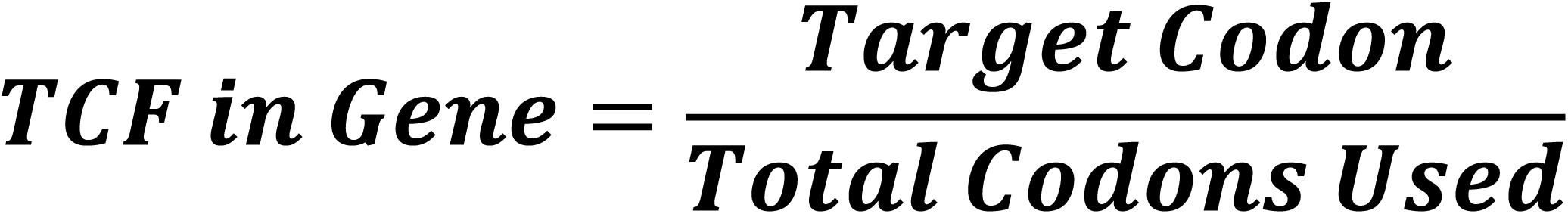

Codon bias analysis of gene groups was performed using T-statistics for both ICF and TCF values. T-statistic calculations use the sample mean (x), the population mean (μ0), the sample standard deviation(s), and the sample size (n) **(Formula 3)**.

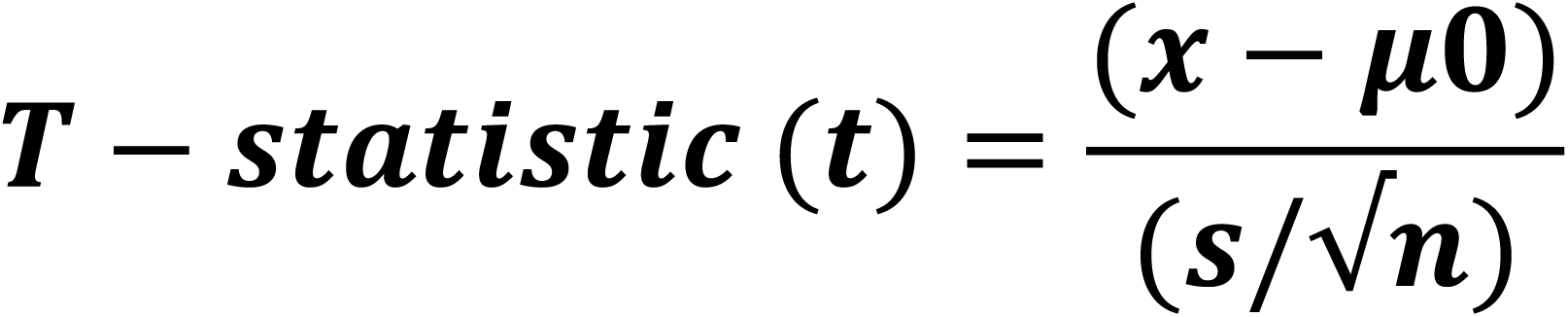

To measure the gene-specific codon bias usage of genes, we used the previously described gene-specific codon usage (GSCU) algorithm (Begley et al., 2007; Chan et al., 2010; et al., 2012; Chionh et al., 2016; Chan et al., 2018). Gene-specific codon bias usage was calculated using Z-scores, which describes standard deviations away from a mean, to indicate whether specific codons were over- or under-represented in a particular gene relative to the genome average usage for that codon **(Formula 4)**.

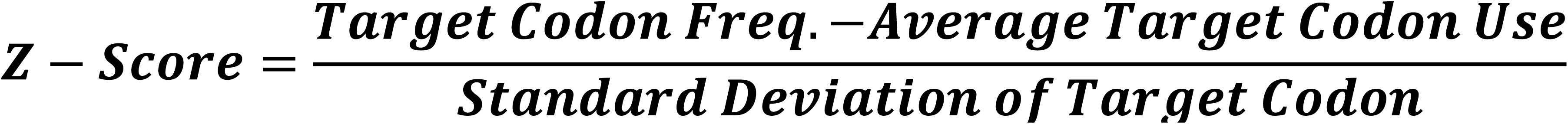

Hierarchical clustering of data and heatmap visualization was performed using Morpheus (https://software.broadinstitute.org/morpheus/) (Broad Institute).

### 2.3 Gene Ontology Enrichment of Oncogenes and TSGs

Diseases associated with oncogene and TSGs sub groups was assessed using gene ontology enrichment analysis by Metascape (https://metascape.org/gp/index.html#/main/step1) (Zhou et al., 2019).

Enrichment cut-offs were set at a p-value < 0.01, a minimum count of 3, and an enrichment factor > 1.5, with the enrichment factor defined as the ratio between observed counts and the number of counts expected by chance.

### 2.4 Gene Ontology and Gene Annotation Analysis

Gene ontology (GO) information and gene annotations were obtained from the AmiGO 2 browser (http://amigo.geneontology.org) (Ashburner et al., 2000; Carbon et al., 2009; Aleksander et al., 2023), which provides access to curated GO term associations across multiple species. The ontology files contain the structured vocabulary of GO terms organized into the three main categories - Biological Process (BP), Molecular Function (MF), and Cellular Component (CC) - along with hierarchical relationships between terms. We specifically looked only at biological processes. The annotation files provide gene-to-GO term associations for specific species. For this study, we downloaded the latest gene annotation (GAF) files for *Homo sapiens* and *Mus musculus* (September 2025), which include mappings of each gene to its associated GO terms. In parallel, we obtained the corresponding ontology (OBO) file to interpret and categorize the annotation data. For each ontology group, we generated gene lists for both *Homo sapiens* and *Mus musculus,* and used them to compare codon usage patterns between oncogenes and TSGs. Next, we integrated codon usage analysis with ontology-based gene classification, thereby linking codon bias signatures to specific biological pathways and functional categories.

### 2.5 PCA Analysis

Principal Component Analysis (PCA) is a dimensionality reduction technique that transforms high-dimensional data into orthogonal principal components (PCs) that capture the maximal variance in the dataset (Jolliffe et al., 2016; Jolliffe et al., 2025). To investigate codon usage preferences across human and mouse oncogenes, we performed PCA using Z-score normalized ICF and TCF values for each gene. PCA results were visualized as scatter plot using ggplot2 in R. Individual genes were plotted according to their PC1 and PC2 coordinates, and colored by their codon usage preference group. Cluster means were calculated for each group, and ellipses representing 95% confidence intervals around these means were drawn using ggforce::geom_ellipse (Pedersen et al., 2024) to highlight the distribution and central tendency of each codon usage group. Similar analyses were performed to compare the codon usage patterns of human and mouse oncogenes. Lastly, PCA was used to compare the codon usage signatures of 447 human GOs and 462 mouse GOs vs both oncogenes and TSGs. We first performed the dimensional reduction on the T-statistics dataset of all GOs followed by Gaussian mixture modelling (GMM) to compare codon usage of oncogenes relative to other GOs (Dempster et al., 1977; Fraley et al., 2002). The top PCs (PC1- PC4) were used as inputs for GMM. We then generated a PCA lattice plot consisting of pairwise scatter plots of PC1 vs PC2, PC1 vs PC3, PC1 vs PC4, PC2 vs PC3, PC2 vs PC4, and PC3 vs PC4. This lattice structure allows simultaneous comparison of multiple principal components to detect codon usage patterns of oncogenes, TSGs, and other 447 human GOs. We also compared mouse GOs by generating PCA and lattice plots in R. For PCA, we used prcomp(), and for lattice visualization, we used pairs() or GGally::ggpairs(). We applied ggplot2 or base graphics for coloring and labeling when needed. A three-dimensional (3D) PCA plot was also generated by plotting sample scores along PC1, PC2, and PC3 axes. This 3D representation allows visualization of multivariate relationships, clustering patterns, and potential separation among GO clusters, oncogenes, and TSGs. This plot was visualized and exported as a two-dimensional (2D) image at a fixed viewing angle, representing a 2D projection of the 3D PCA space.

## 3. Results

### 3.1 Human oncogenes have a distinct codon usage bias relative to TSGs and other groups of genes

To identify codon usage signatures distinct to oncogenes and TSGs, we first compared individual codon frequencies between each class of genes by computing T-statistics based on either ICF or TCF. A positive T-statistic identifies a group of genes with values higher than the genome average (HTA) for a specific codon, with negative values signify lower than average (LTA) codon usage. Using ICF values, we determined 8 overlapping (HTA vs. HTA or LTA vs. LTA) and 54 opposing (HTA vs. LTA or LTA vs. HTA) codons for oncogenes vs. TSGs **(Figure 1A)**. As a group, oncogenes overuse codons that end in G/C (GC3), while TSGs favor codons that end in A/U (AU3). We also observed 17 overlapping and 45 opposing codon trends when comparing human oncogenes vs TSGs using TCF values **(Figure 1B)**, with similar trends in GC3 and AU3 observed for TCF-based analysis. Heat map visualization and hierarchical clustering (based on Euclidian distance) of T-statistics also showed similar trends and highlight opposing codon usage patterns of oncogene vs. TSGs using both ICF- and TCF-based metrics **(Figure S1, Supplemental Tables S1A and S2A)**. ICF values indicate usage biases in synonymous codons coding for the same amino acid, while TCF values indicate overall codon usage biases in the gene and may be skewed due to the gene’s amino acid composition. The ICF and TCF data with T-statistics, p-values, codon counts, and Z-score data used to generate the bar graphs and heatmap are detailed in **Supplemental Tables S1-4.** Importantly, these patterns were recapitulated when the analysis was repeated using an independently curated list of oncogenes and TSGs. When using the ICF matrix, for this second list of genes, we found 12 overlapping and 50 opposing codon trends statistics for oncogenes vs TSGs. Similarly, the TCF metric showed 17 overlapping and 45 opposing codon trends **(Figure S2, Supplemental tables S1B and S2B).** The concordance between datasets supports the robustness and reproducibility of the identified opposing codon usage signatures between human oncogenes and TSGs.

**Figure 1.**
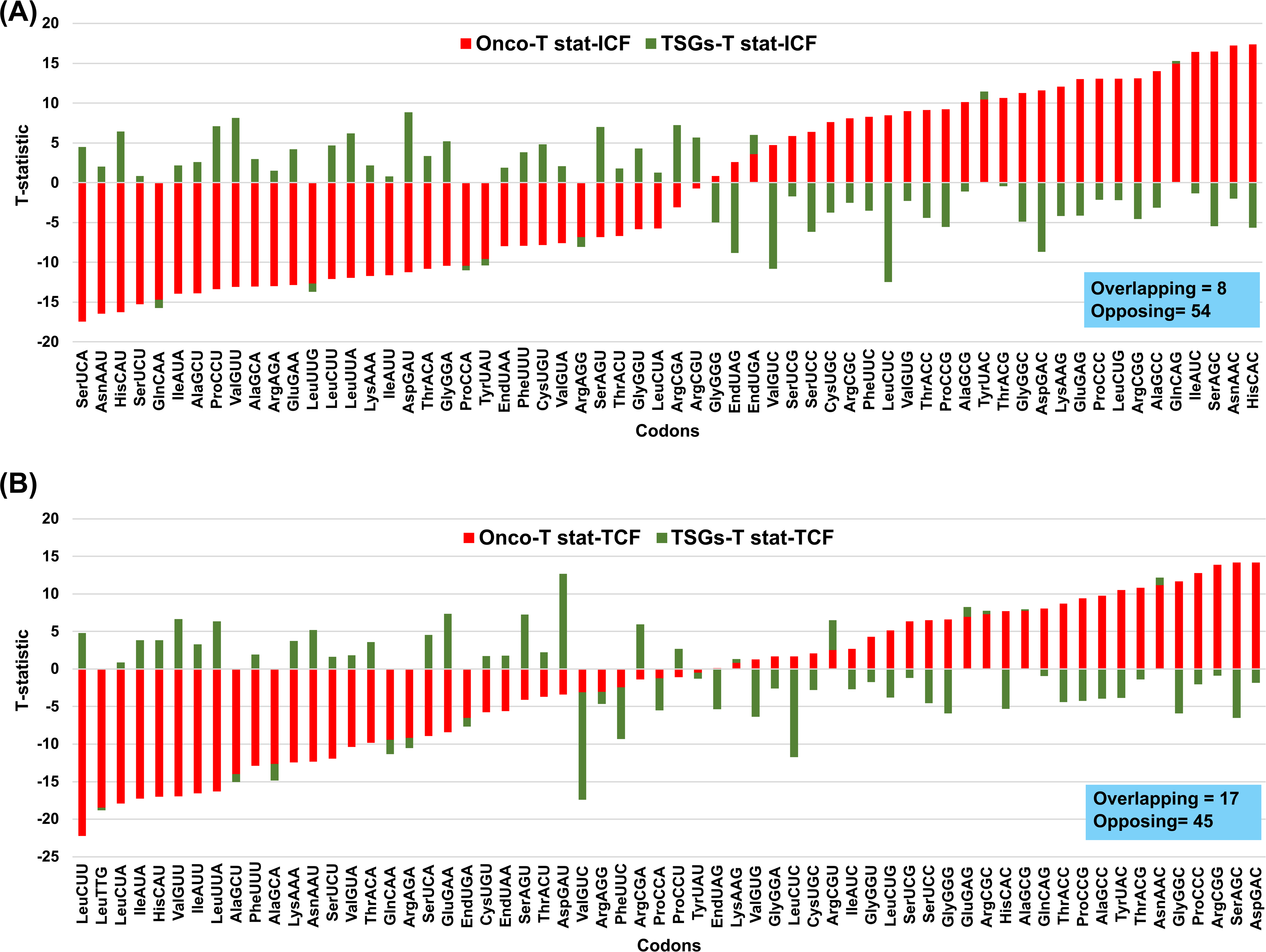
Human oncogenes have distinct codon usage patterns relative to human TSGs. Stacked bar plots showing comparative trends in codon usage bias between human oncogenes and TSGs using T-statistics for ICF and TCF metrics. Red bars represent human oncogenes and green bars represent human TSGs. Codons are along the x-axis and T-statistics on the y-axis, where positive and negative value identifies a group of genes that have values higher or lower than average values, respectively. Overlapping bars indicate the number of codons where the T-statistics are in same direction for both oncogenes and TSGs, while opposing bars indicate codon usage in opposite directions. The number of codons used in respective directions is in blue box. **(A)** ICF- and **(B)** TCF-based T-statistics.

### 3.2 Most oncogenes are GC3 biased, with a small group trending to AU3 biased

Gene-specific codon usage (GSCU) Z-score values detail whether a gene over- or under-uses a codon relative to genome average usage of that codon. To determine codon biases within each specific gene, we computed codon specific Z-scores for 66 human oncogenes using ICF or TCF data **(Supplemental Tables S3 and S4).** Using hierarchical clustering of ICF linked Z-scores **(Figure 2A)**, we determined that human oncogenes have three major groups of genes: (i) 43 genes over-use G/C-ending (GC3) codons, (ii) 12 over-use A/U-ending (AU3) codons, and (iii) 11 genes with codon usage that resembles genome averages. Specific to TCF, there are (i) 42 genes that over-use G/C-ending codons, (ii) 12 genes that over-use A/U-ending codons, and (iii) 12 genes that resemble genome averages **(Figure 2B)**. Principal component analysis (PCA) reinforces these findings regarding gene-specific codon usage clusters within the oncogene group (**Figure 2C)**. On the whole, oncogenes are strongly biased towards GC3 codons, with some showing an extreme preference (e.g., FGF4, CCND1, SKI, etc.), while fewer over-use AU3 codons (e.g., MDM2, KRAS, REL, etc) as shown in **Figure S3A.** Human tumor suppressor genes also have one small group which over-use GC3 codons (21 genes) with some extremes such as BRCA2, PMS1, PTEN, etc. Another larger group shows an extreme preference for AU3 ending (33 genes), with CEBPA, BCL11B, NOTCH1 being some of the most pronounced examples, and still others with no bias at all. (**Figure S3B).** Next, gene-specific average Z-scores and frequencies of human oncogenes and TSGs were determined by calculating the average of the Z-score values for each codon in a group (all genes) and subgroup (genes over-using GC3/AU3 codons), which acts as a measure of codon bias. We found opposite trends in human oncogenes and TSGs when looking at Z-score values, but did not notice many differences in codon usage when comparing their frequencies alone **(Supplementary Table S5).** The Z-score data for human oncogenes, can be found in **Supplementary Tables S6A (ICF)** and **S6B (TCF)**, and for human oncogenes frequency data, refer to **Supplementary Tables S6C (ICF)** and **S6D (TCF).** For human TSG average Z-score data, refer to **Supplementary Tables S7A (ICF)** and **S7B (TCF)**, and for human TSG frequency data, refer to **Supplementary Tables S7C (ICF)** and **S7D (TCF).**

**Figure 2.**
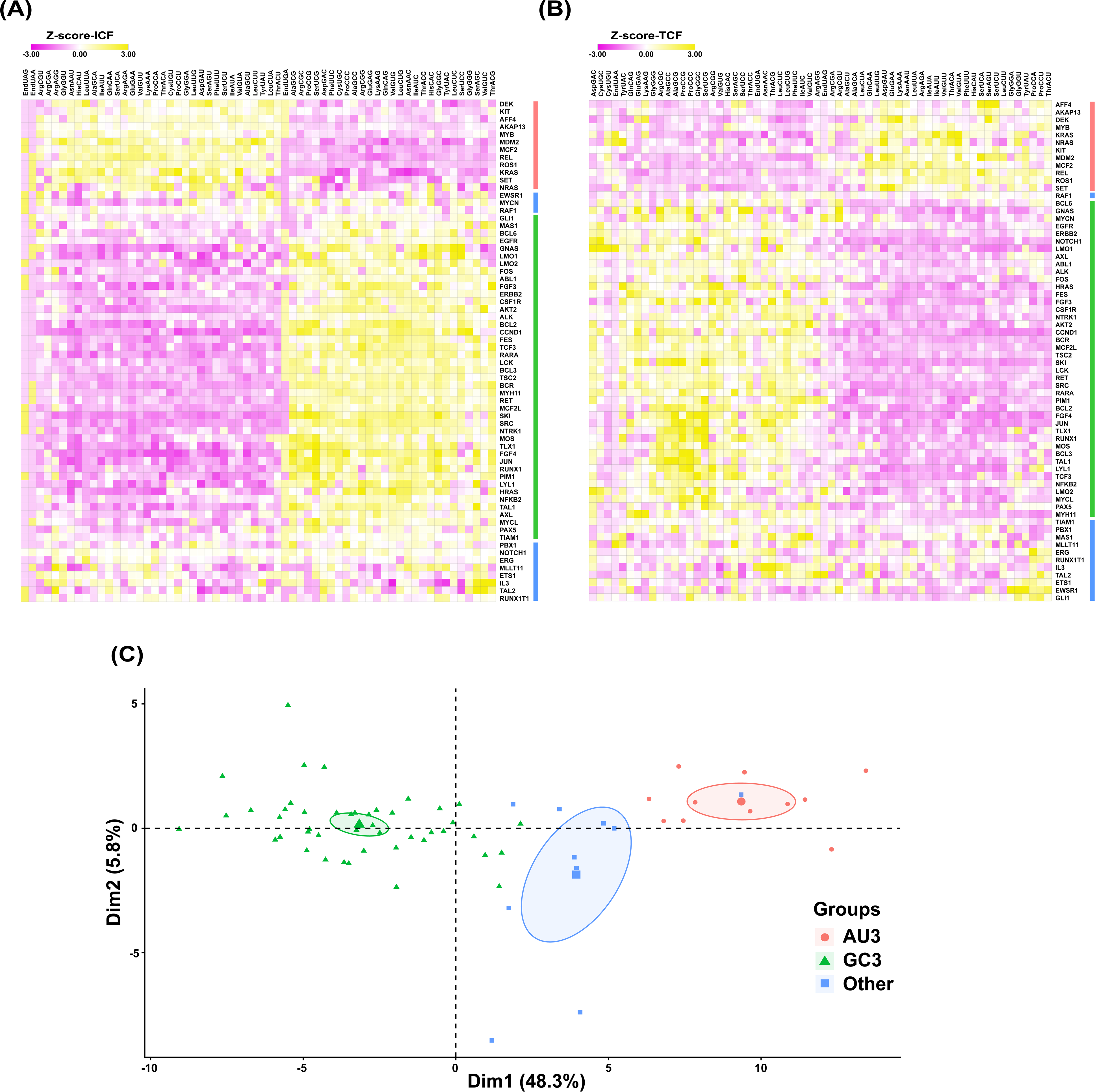
Extreme codon usage bias signatures in human oncogenes. ICF and TCF Z-scores of human oncogenes were hierarchically clustered and visualized as heatmap, describing the gene-specific codon usage (GSCU) for each human oncogene, relative to the genome average **(A)** ICF and **(B)** TCF. Rows represent oncogenes and columns represent codons. Color intensity indicates Z-score values depicting the over-use or under-use of codons relative to the genome average. Green, red and blue bars next to heatmaps show group of genes enriched in GC3, AU3 and no change respectively. **(C)** Principal component analysis (PCA) of codon usage patterns in human oncogenes using ICF Z-scores. Each point represents an oncogene, colored by its codon usage preference. Genes preferring AU-ending codons (red), GC-ending codons (green), and other genes with no change (blue).

### 3.3 GC3 and AU3 biased oncogenes can be linked to specific cancers

To identify ontologies and specific cancers linked to the oncogenes that over-use GC3 (N = 43) or AU3 (N = 12) codons according to ICF, we used Metascape (http://metascape.org/gp/index.html#/main/step1) analysis. Oncogenes over-using GC3 codons were enriched in GO terms including congenital chromosomal disease, leukemia, kidney Wilm’s tumor, and gastrointestinal stromal tumors **(Figure 3A).** Oncogenes over-using AU3 codons were significantly enriched in myeloproliferative disease, congenital chromosomal disease, and leukemia, among others **(Figure 3B).** A heatmap based on -log10 (p values) showed that the oncogenes over-using AU3 codons are uniquely enriched in T-cell lymphoma, childhood kidney Wilm’s tumor, malignant neoplasm of soft tissue, nephroblastoma, lymphoid leukemia, acute leukemia, B-cell malignancy, gastrointestinal stromal tumors, mammary neoplasms, and mammary carcinoma. On other hand, oncogenes over-using GC3 codons have unique enrichments such as myeloproliferative disease, blast phase, papilloma, HER2 gene amplification, follicular lymphoma, papillary carcinoma, and various kinds of leukemia **(Figure 3C).**

**Figure 3.**
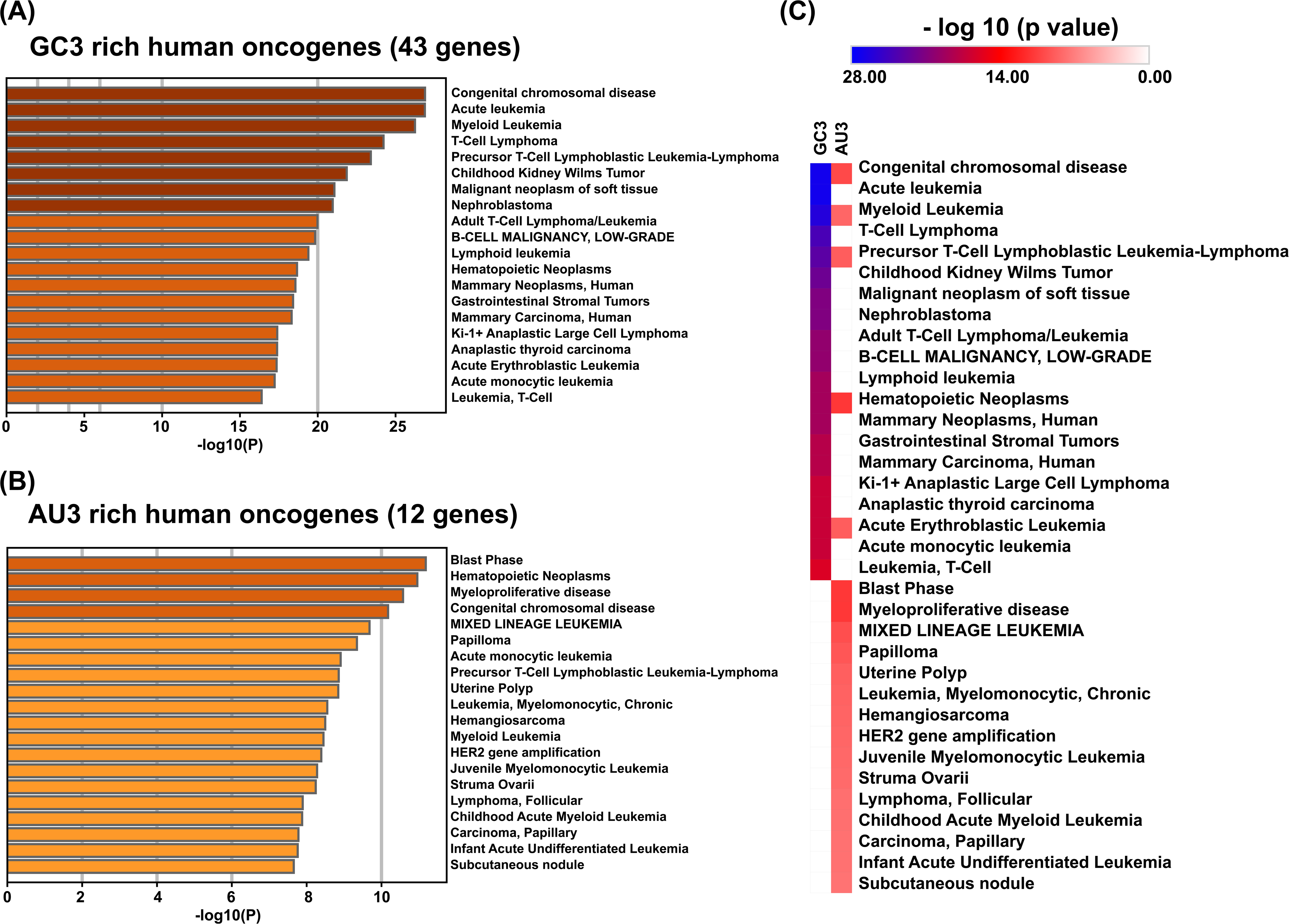
Cancer types can be linked to GC3 and AU3 codon biased oncogenes. Gene ontology and disease enrichment analysis of human oncogenes stratified by codon usage bias. Human oncogenes have three groups of genes according to their codon usage preference: genes preferring GC-ending codons, genes preferring AU-ending codons, and genes with no change in their codon usage. **(A)** Enrichment analysis of human oncogenes preferentially using G/C-ending codons, identifying significantly associated cancer types and disease terms. (B) Enrichment analysis of oncogenes preferentially using A/U-ending codons, highlighting disease associations distinct from or shared with the G/C-biased group. (C) Heatmap summarizing overlapping and uniquely enriched disease terms between A/U- and G/C-codon–biased oncogenes using their log_10_- transformed p-values. Color intensity reflects enrichment significance, allowing comparison of shared and group-specific disease associations. Gene ontology and disease enrichment analyses were performed using Metascape.

### 3.4 Mouse oncogenes also display distinct codon usage biases when compared to TSGs

Similar to the analysis performed for human oncogenes, we used group-based measures of codon usage to compare mouse oncogenes and TSGs to determine whether the observed codon usage patterns for humans can be generalized across species **(Figure 4, Supplemental Tables S8 and S9)**. Using ICF-based values, we identified 25 overlapping and 37 opposing codon trends **(Figure 4A)**, while for TCF-based values we observed 34 overlapping and 28 opposing codon trends (**Figure 4B).** We also used heat map-based visualization of hierarchically clustered ICF and TCF linked T-statistics to compare mouse oncogenes and TSGs **(Figure S4.A – B, Supplemental Tables S8 and S9)**. These results show that, similar to observations in humans, mouse oncogenes have distinct codon usage patterns relative to mouse TSGs. In addition, when T-statistic values for oncogenes and TSGs for humans and mice groups were compared, we found that humans have more extreme differences compared to mice **(Figure S4C, Supplemental Table S12).** ICF- and TCF-based Z-scores for individual mouse oncogenes were also determined **(Figure. 5, Supplemental Tables S10 and S11)**. Similar to humans, there are three major groups of mouse oncogenes, ones over-using either AU3 or GC3 codons, and a group with little codon bias. For ICF, there are 23 mouse oncogenes over-using GC3 codons, 14 genes over-using AU3 codons, and 8 genes close to the genome average **(Figure 5A)**. For TCF, there are 20 mouse oncogenes over-using GC3 codons, 13 over-using AU3 codons, and 12 genes showing little codon bias **(Figure 5B)**.

**Figure 4.**
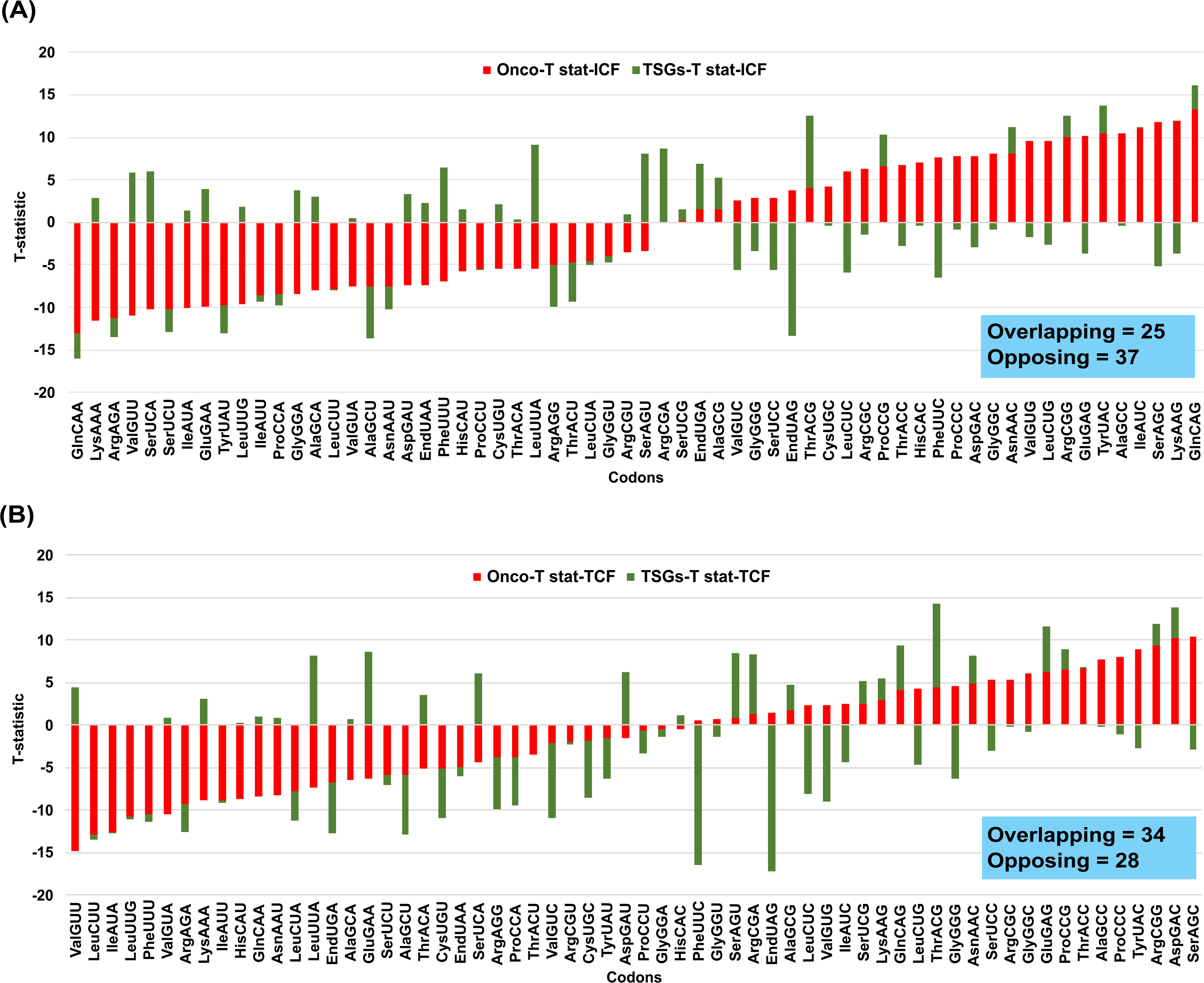
Mouse oncogenes have distinct codon usage patterns relative to mouse tumor suppressor genes. Stacked bar plots showing comparative trends in codon usage bias between mouse oncogenes and tumor suppressors using T-statistics for ICF and TCF metrics. Red bars represent mouse oncogenes and green bars represent tumor suppressors. Codons are along the x-axis and T-statistics on the y-axis, where positive and negative value identifies a group of genes that have values higher or lower than average respectively for a specific codon. Overlapping bars indicate the number of codons where the T-statistics are in same direction for both oncogenes and tumor suppressors, while opposing bars indicate diverging codon usage bias. The number of codons used in each respective direction is in blue box. **(A)** ICF and **(B)** TCF.

**Figure 5.**
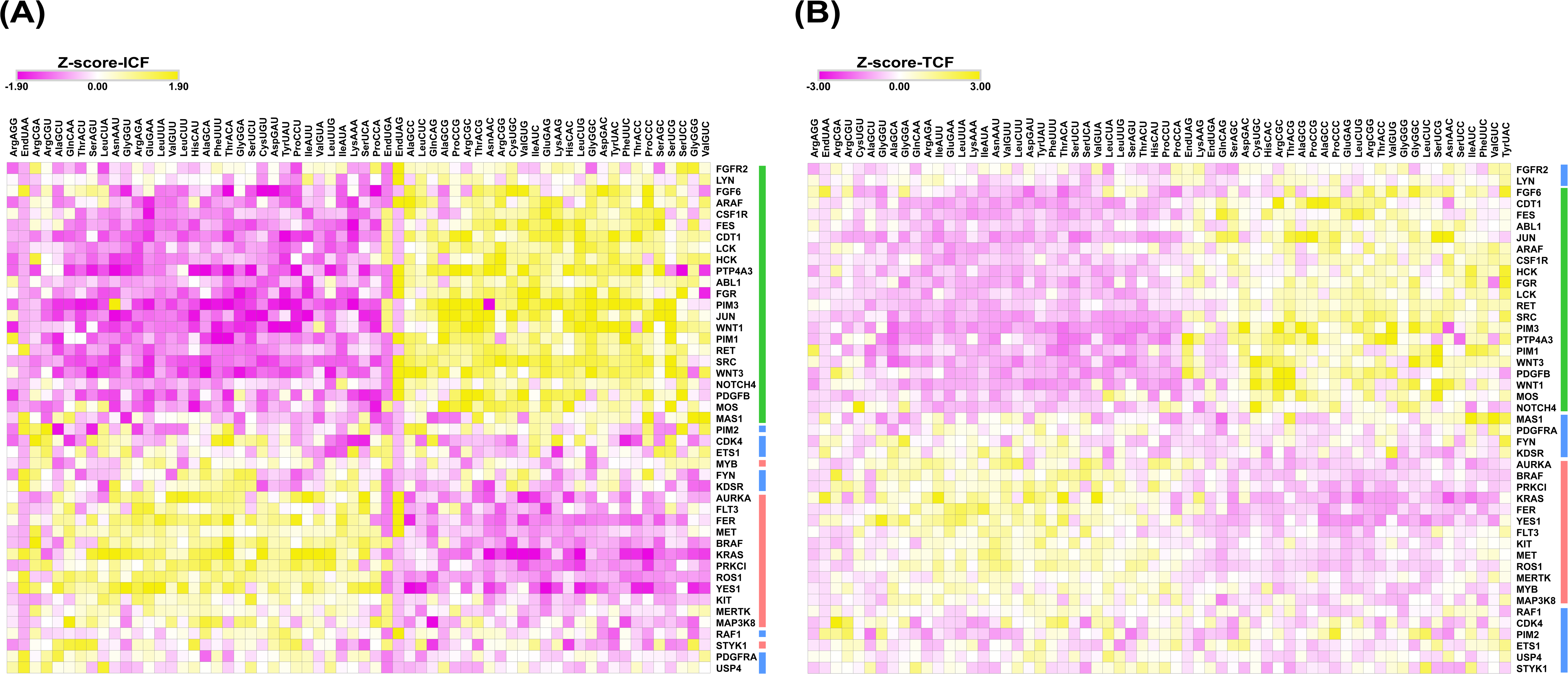
AU3 or GC3 codon bias in mouse oncogenes. Z-scores of mouse oncogenes were hierarchically clustered and visualized as heatmap, describing the gene-specific codon usage (GSCU) for each mouse oncogene relative to the genome average (A) ICF and (B) TCF. Rows represent oncogenes, and columns represent codons. Color intensity indicates Z-score values depicting over-use or under-use of codons relative to genome average. Green, red and blue bars next to heatmaps show group of genes enriched in GC3, AU3, and other genes with no change relative to genome average respectively.

### 3.5 Codon bias usage signatures of oncogenes are distinct compared to other gene ontologies

We used AmiGO2 to obtain gene ontology (GO) annotations for human and mouse species to systematically examine codon usage biases within functional gene groups to determine whether codon bias patterns are distinct to oncogenes and TSGs. We compared the T-statistics of both ICF- and TCF-based measures for 447 human gene ontology-based groupings, all with between 50 to 150 genes, to oncogenes and TSGs **(Supplemental Table S13).** A heatmap of T-statistics for ICF-based measures revealed clear and distinct codon usage signatures for oncogenes relative to the other 447 GOs and TSGs **(Figure 6A)**. To further investigate codon-based patterns in GOs, we performed PCA using the ICF-based T-statistic measures and visualized the top four principal components (PC1–PC4) using a lattice of pairwise scatter plots. Together, these components captured the majority of variance in the dataset. We found six clusters in the data. The PCA lattice showed that oncogenes and TSGs appeared as separate points along PC1 and PC2, which explained 70% and 6.2% of the variance respectively. This pattern reflects differences in their codon usage profiles **(Figure S5A).** PC3 and PC4 explained much less variance, at 3.2% and 2.4%, and mainly captured differences within groups, especially among GO categories, without further separating oncogenes and TSGs. To further interpret the principal component analysis, mean PCA scores for the first four principal components (PC1–PC4) were compared among Gene Ontology (GO) genes, oncogenes, and TSGs. The bar graphs summarize the average position of each group along the major axes of variance captured by the PCA. Distinct differences in mean scores were observed across groups, indicating that GOs, oncogenes, and TSGs contribute differently to the variance represented by the principal components **(Figure S5B).** To visualize the multivariate relationships among the three groups, a three-dimensional (3D) PCA plot based on PC1, PC2, and PC3 was generated **(Figure 6B)**. For presentation purposes, the 3D PCA space was displayed and exported as a two-dimensional (2D) projection at a fixed viewing angle, allowing simultaneous visualization of the first three principal components. The projected PCA plot revealed clear spatial organization for 6 clusters of GO terms, oncogenes, and TSGs within the reduced dimensional space. We calculated average T-statistic values for GC3 and AU3 codons in each cluster and compared oncogenes and TSGs, with this data clearly showing that oncogenes occupy a significantly distinct codon space **(Figure 6C).** We observed similar trends in human TCF-based data shown **(Figure S6 and S7).** A similar analysis was performed on 462 mouse gene ontology–based gene groups, each comprising approximately 50–150 genes, using their T-statistics for both ICF- and TCF-based metrics **(Supplemental Table S14).** The analysis in mice revealed structured but less extreme separation between oncogenes, TSGs, and other GO’s, when compared to humans **(Figure S8 – S9)**.

**Figure 6.**
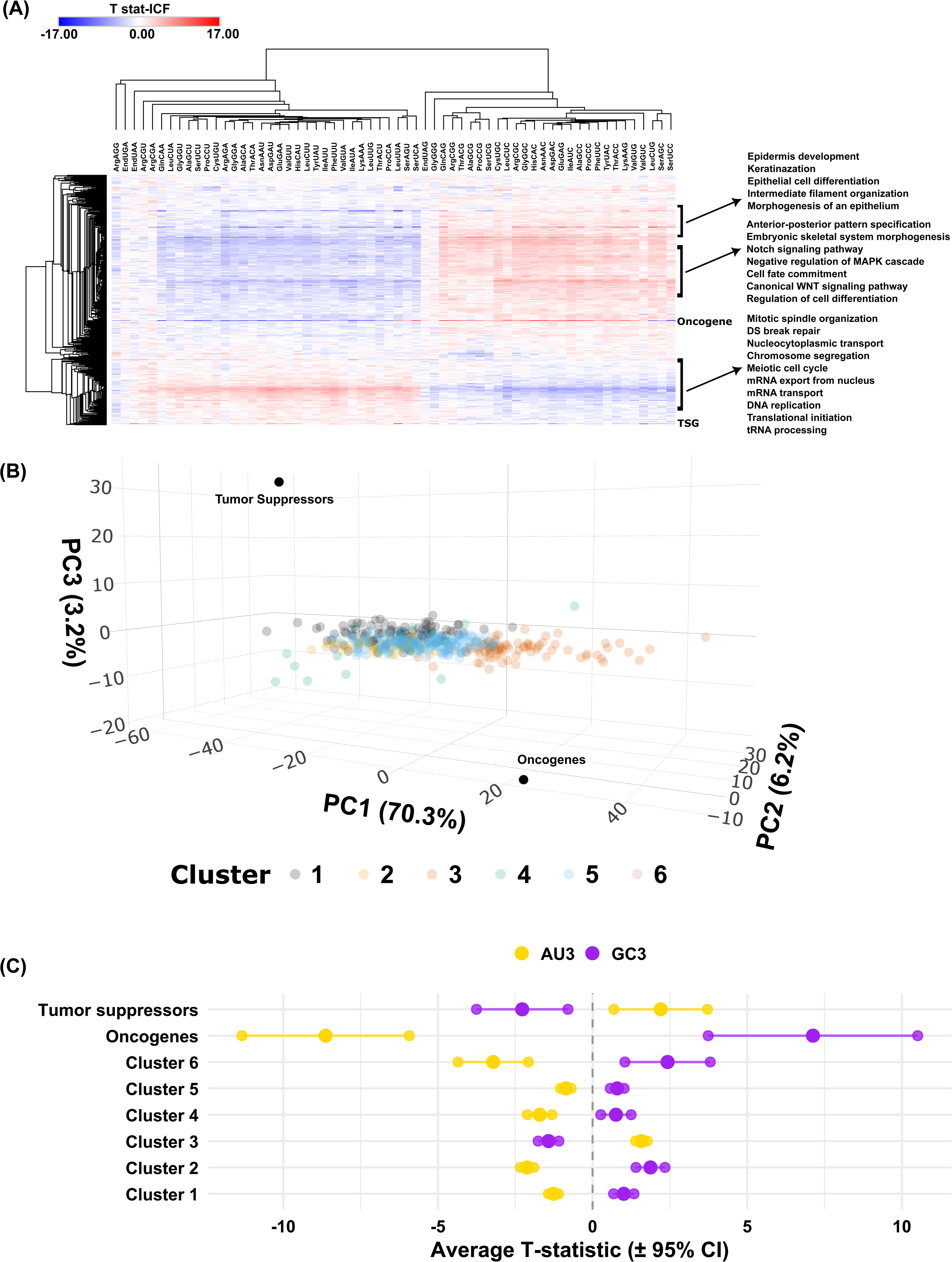
Gene Ontology (GO) based gene annotation of human oncogene and tumor suppressors. Heatmap of **(A)** ICF t-statistic values for 447 GOs, human oncogenes, and human tumor suppressor genes. They were hierarchically clustered, describing codon usage patterns of all categories. **(B)** A 3D PCA score plot was generated using the first three principal components (PC1, PC2, and PC3) and visualized as a 2D projection at a fixed viewing angle. Each dot represents an individual GO term. Human oncogenes and tumor suppressors are labelled as black dots. **(C)** The cluster map shows AU3 and GC3 bias across six clusters of human GOs, oncogenes, and TSGs. The values represent T-statistics, where positive values indicate enrichment and negative values indicate depletion. Bias was considered significant when 95% CIs did not overlap.

## Discussion

### Codon bias is an organizational code

Codon usage patterns have previously been used to stratify stress response and other proteins, with distinct patterns of codon bias linked to translational regulation by specific tRNA-based systems (Endres et al., 2015b; Hanson et al., 2018; Mitchener et al., 2023; Wu et al., 2024). For example, stress-responsive transcripts can be selectively translated through tRNA modification–dependent mechanisms (so-called modification-tunable transcripts), where changes in wobble-uridine tRNA modifications alter the decoding efficiency of specific synonymous codons enriched in certain classes of mRNAs (Begley et al., 2007; Dedon et al., 2014). Our findings highlight that codon usage bias can function as an “organizational code” for biologically related gene groups, which could potentially bin them into protein expression potential and outputs. Synonymous codon usage provides translational choices and can contribute regulatory elements to gene expression, especially in genes requiring tight control, such as oncogenic programs. For instance, rare synonymous codons regulate KRAS translation and protein levels (Lampson et al., 2013; Fu et al., 2018), and synonymous mutations can act as cancer drivers by altering translational regulation (Supek et al., 2014; Li et al., 2021). Synonymous codon composition (optimal vs non-optimal codons) can affect translation initiation, with non-optimal codons leading to reduced engagement by ribosomes and translational repression that is not solely explained by mRNA levels (Barrington et al., 2023). Human oncogenes occupy an extreme and unique codon-usage space in the genome which may translationally pre-program corresponding mRNAs for rapid, robust, or condition-dependent expression. Further, our findings comparing 447 GO-categories to oncogenes and TSGs supports the idea that functionally similar genes have distinct patterns based on codon usage (Sharp et al., 1987; Plotkin et al., 2011; Gingold et al., 2014; Pershing et al., 2015; Presnyak et al., 2015; Ghanegolmohammadi et al., 2026). Previous studies have demonstrated that simple measures of codon usage bias can define DNA damage response, ROS-detoxification, and translation linked genes, among others (Begley et al., 2002; Begley et al., 2007; Chan et al., 2010; Gu et al., 2014), with advanced correlation analysis recently using codon usage signatures to identify 91 gene families in yeast (Ghanegolmohammadi et al., 2026).

### Contrasting codon bias has systems level regulatory potential

Distinct codon bias may act as a systems-level feature allowing cells to regulate the protein synthesis machinery tell oncogenes and tumor suppressor genes apart from mRNAs involved in other biological processes, with acceleration of oncogenes. This can speed up oncogene activity and slow down tumor suppressor genes, which can drive cell proliferation programs. Functionally, oncogenes broadly act to promote cellular growth and proliferation. For example, growth-factor signaling oncogenes such as RAS activate signaling that drive cell-cycle progression, survival, and metabolic reprogramming in diverse cancer types (Pylayeva-Gupta et al., 2011; Yang et al., 2024). Oncogenes such as MYC promotes cell growth by stimulating ribosome biogenesis, metabolism, and cell-cycle progression (Dang et al., 2012; Seres et al., 2025). In addition, the receptor and kinase oncogenes such as EGFR and BRAF enhance mitogenic signaling and uncontrolled division when aberrantly activated (Shan et al., 2024; Seres et al., 2025). In contrast, tumor suppressor genes act to restrain growth and maintain cellular checkpoints. Loss or mutation of the TSG TP53 undermines DNA-damage responses and apoptotic programs that normally limit aberrant growth, with recent work showing that even synonymous mutations in TP53 can alter isoform expression with potential functional consequences for tumor suppression (Levine et al., 2009; Sajwan et al., 2025). Other TSGs such as RB1 and PTEN maintain cell-cycle checkpoints and inhibit pro-growth signaling, with dysregulation leading to unchecked proliferation (Dick et al., 2013; Stojchevski et al., 2025), signaling to promote survival and proliferation (Song et al., 2012; Stojchevski et al., 2025). Together, these opposing functional classes illustrate how growth-promoting versus growth-restricting programs operate at a systems level, consistent with the idea that their transcripts may also be distinguished by regulatory features such as codon usage. The observation that human oncogenes and TSGs have opposing codon biases supports the idea that there could be opposing regulatory effects **(Figure 7).** Expression programs that favor the translation of GC3 rich oncogenes could correspondingly inhibit the synthesis of AU3-rich TSGs and act as a programmatic trigger to accelerate proliferation via increased oncogene levels, while removing TSG-based brakes that slow down cell growth. This on-off regulatory relationship between oncogenes and TSGs has been reported for ploidy changes and mutations (Beroukhim et al., 2010), with breast cancer findings of amplified ERB2/HER2 (GC3) and inactivated p53 (AU3) (Network et al., 2012; Levine et al., 2013), as well as pancreatic cancers with activating mutations in KRAS (AU3) and corresponding mutations or deletions of BRCA1, BRCA2, or ATM (all GC3) (Varghese et al., 2025).

**Figure 7.**
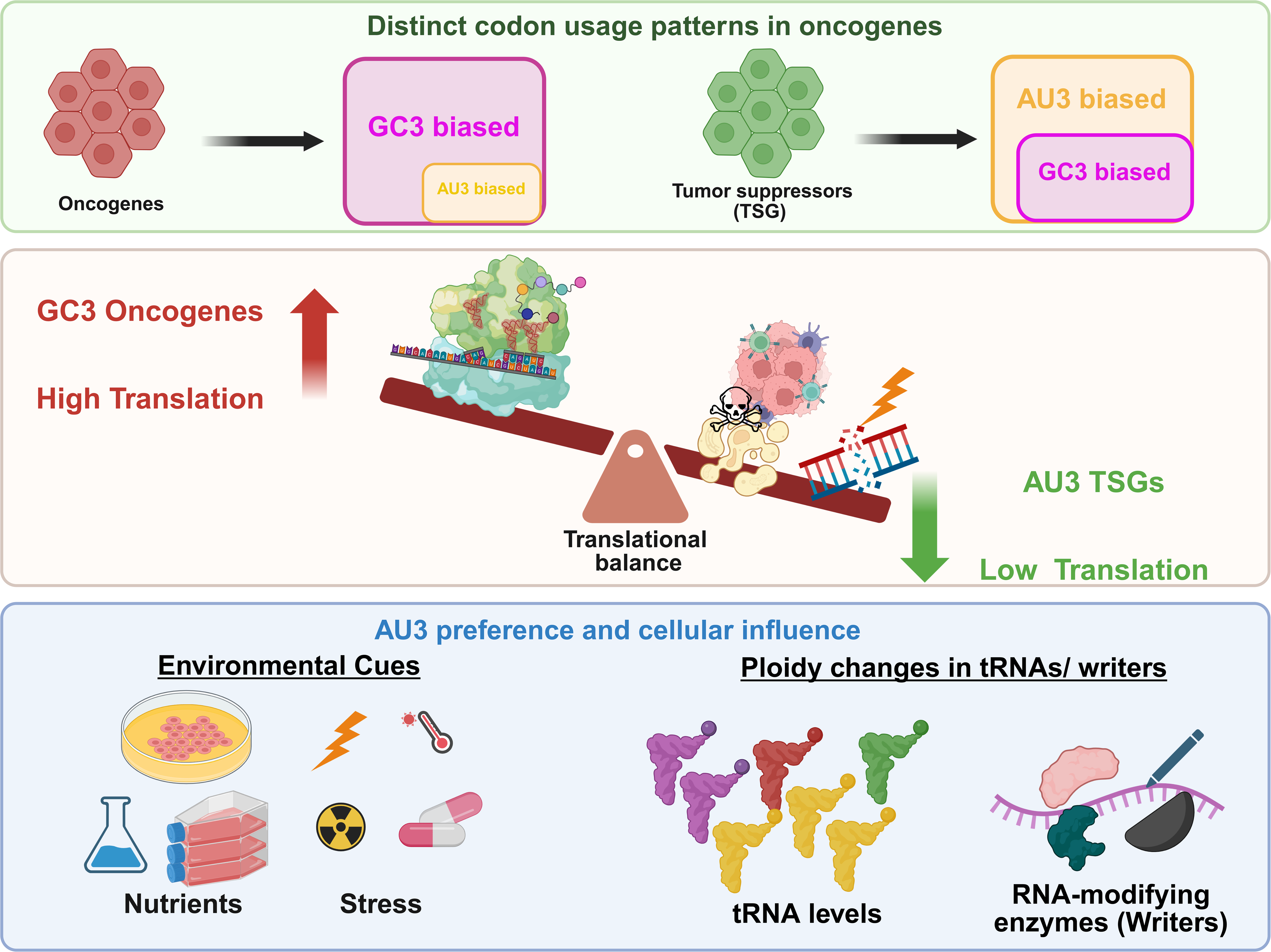
Schematic representation of codon usage bias–driven translational regulation of oncogenes. Codon bias features and contrasts in oncogenes and TSGs (top) and potential mechanism of systems level control to promote cell proliferation (middle). A switch to AU3 based translation can occur by environmental or genetic changes (bottom).

### Cells favor and have potential to change translated codon bias

The extreme codon biases observed in some oncogenes support evolutionary models proposing that cancer-related genes are optimized for rapid or conditional expression (Sharp et al., 1987; Pavon-Eternod et al., 2009; Gingold et al., 2014; Goodarzi et al., 2016). Gingold et al. (2014) showed that proliferating and differentiating cells use different tRNA pools that match the codon usage of genes active in each cell state. This means the translational machinery can shift to favor certain transcripts depending on the cell’s context. Building on this, our results show that oncogenes and TSGs also sit at opposite ends of the codon usage spectrum. GC3-enriched oncogenes are linked to translational programs seen in proliferating cells, while AU3-enriched TSGs may match tRNA pools found in growth-restricted states. These findings suggest that cancer cells use tRNA reprogramming machinery, which usually helps cells switch from proliferation to differentiation, to boost oncogene translation and reduce TSG output. This adds a new layer to how gene expression is controlled in cancer (Gingold et al., 2014; Goodarzi et al., 2016). We recently demonstrated that an extreme natural codon bias in the *CEBPB* gene for 4 C-ending codons for Tyr, His, Asn, and Asp, which aligns with a GC3 bias, promotes dramatically increased protein levels of CEBPB protein relative to an engineered version that switch to the U ending version (Davis et al., 2025). Similarly, extreme AU3 bias found in MIER1 and corresponding low expression levels in HEPG2 cells was rescued by codon re-engineering to C-ending codons for Tyr, His, Asn, and Asp. These findings align with our current study and highlight that codon bias has regulatory effects. Further it suggests that HEPG2 cells are optimized to translate GC3 ending codons (Davis et al., 2025). The AU3-biased oncogenes we found suggest that certain cells may need to be adjusted or encouraged to translate AU3 enriched codons more effectively. AU3-biased oncogenes may represent a stress-responsive translational subgroup and suggest alternative translational programs could be used to optimize their expression, relative to GC3. Stress, environmental cues, or specific tRNAs and corresponding RNA modifications could act as triggers for AU3 translation, and this process could give AU3 routes new ways to increase oncogene expression **(Figure 7)**. The potential of an AU3 translational program suggests that some oncogenes may have enhanced expression under specific environmental stress or altered tRNA states. For example, METTL1 based expression of AU3-linked oncogenic programs has been linked to writer amplification via the stabilization and enhanced levels of specific tRNAs (Lin et al., 2018; Orellana et al., 2021; Li et al., 2023). Similarly, overexpression of ELP3, which is essential to the formation of mcm^5^-based modifications in tRNA^(UUU)^_Lys_, tRNA^(UUG)^ , and tRNA^(UUC)^ has also been shown to drive cancer proliferation and chemotherapeutic resistance, with ELP changing translational programs (Ladang et al., 2015; Delaunay et al., 2016). In addition to alterations in tRNA writers, stress and changing physiological conditions have been shown to re-program a spectrum of tRNA modifications, which can alter the translation and levels of key codon biased genes (Chan et al., 2012; Begley et al., 2013; Dedon et al., 2022; Huber et al., 2022). Notably, stress from reactive oxygen species (ROS) has repeatedly been shown to alter the epitranscriptome in multiple systems, with altered ROS maintenance and level linked to multiple cancers (Chan et al., 2012; Endres et al., 2015a; Zhang et al., 2016; Leonardi et al., 2019; Huber et al., 2022).

### Mice have less extreme codon bias than humans

Mouse codon bias specific to oncogenes and TSGs is present but clearly not as dramatic as that observed in humans. Species-specific codon programs may limit translation-based extrapolation from mouse cancer models to humans (Rangarajan et al., 2003; De Jong et al., 2010; Ellis et al., 2010), with the difference in codon bias highlighting considerations for experimental design and interpretation of findings from some for mouse cancer studies. Consistent with a translational control framework, numerous mouse oncogene studies show that growth promoting signaling pathways actively reprogram protein synthesis capacity and selectivity. In MYC-driven mouse tumors, MYC activation is associated with increased ribosome production, elevated tRNA and rRNA synthesis, and broadly enhanced cap-dependent translation, collectively expanding translational output and tumor growth (Barna et al., 2008; Ruggero et al., 2009; Van Riggelen et al., 2010). Similarly, constitutive activation of the PI3K–AKT–mTOR pathway in mouse models promotes cell growth by stimulating translation initiation via mTORC1, 4E-BP phosphorylation, and S6 kinase signaling, resulting in greater ribosome engagement and increased protein production, with many tumors showing continued reliance on these translational control nodes (Thoreen et al., 2009; Hsieh et al., 2012; Thoreen et al., 2012). In parallel, RAC1-driven oncogenic programs in mice have been linked to enhanced proliferation and invasiveness, in part through mTOR-connected translational regulation and coordination between cytoskeletal dynamics and protein synthesis (Kissil et al., 2007; Saci et al., 2011). Taken together, these findings support the idea that multiple mouse oncogenes contribute to tumor development not only through transcriptional and signaling effects, but also by directly elevating translational throughput, reinforcing the relevance of codon-dependent and tRNA-dependent regulatory layers. Our findings suggest that species-specific codon usage differences may shape how these translational programs are implemented across organisms. A unifying principle in mouse and human cancer biology, is that both oncogene activation and tumor suppressor loss converge on control of translational capacity, ribosome biogenesis, and selective mRNA recruitment. Within this framework, codon- and tRNA-dependent regulatory layers represent a mechanistic substrate upon which oncogenic signaling programs operate. However, given that mouse codon bias landscapes appear less polarized than those in humans, species-specific codon usage differences may influence translational efficiency, mRNA selectivity, and therapeutic vulnerability. Thus, while murine models robustly demonstrate that tumorigenesis is tightly coupled to reprogramming of protein synthesis, interspecies variation in codon architecture and translational control environments should be carefully considered when extrapolating translation-centric findings from mouse systems to human cancers.

### Therapeutic implications of oncogene and TSG codon bias

Studies have shown that selective depletion of specific tRNAs via RNA interference impairs translation of mRNAs enriched for matching codons linked to metastasis-associated transcripts, directly linking tRNA abundance to oncogenic protein output (Goodarzi et al., 2016; Orellana et al., 2021). These findings suggest that altering tRNA availability rewires translation of codon-defined gene networks. Our codon analytics findings highlight extreme codon bias in oncogenes that is contrasted by an opposite bias in TSGs. Exploiting extreme codon usage bias represents a potential target for therapeutics that decrease or prevent the translation of specific oncogenes. Therapies that alter codon usage patterns, target specific tRNAs, or inhibit particular RNA modifications have the potential to decrease the expression of extremely GC3-rich oncogenes (i.e., CCND1, FG4, and SKI) or AU3 oncogenes (i.e., KRAS, MDM2, and REL). In addition, being able to selectively change a tumor’s translation from GC3-promoting and AU3-restricting to GC3-restricting and AU3-promoting (and vice versa) could provide a unique way to slow cell proliferation in specific cancers. Yet to be developed therapeutics that specially altered oncogene translation based on codon usage may affect translation elongation, which could alter speed and fidelity. Disrupting translational fidelity has precedence as the cornerstone of antibiotic use to treat infections, and altering translation and fidelity in cancer cells has great potential. Targeting specific codon–tRNA interactions linked to codon bias could enable a very selective targeting approach for specific cancers, and work in this area has the potential to exploit a regulatory feature of cancer programs.

## Data availability statement

The original contributions presented in the study are included in the article/Supplementary Material, further inquiries can be directed to corresponding author.

## Author contributions

CM: Conceptualization, Data curation, Formal Analysis, Investigation, Methodology, Validation, Visualization, Writing–original draft, Writing–editing. ED: Conceptualization, Formal Analysis, Investigation, Validation. DE: Writing-editing, Data curation, Validation, Visualization, Formal analysis. HO: Visualization, editing. SB: Writing–review and editing, Visualization. UB: Methodology, Visualization. LE: Writing–review and editing. PCD: Conceptualization, Supervision, Validation, Writing–review and editing. TJB: Conceptualization, Formal Analysis, Supervision, Validation, Visualization, Writing–review and editing, resources, funding acquisition.

## Funding/ acknowledgements

The authors are grateful for support from the National Institutes of Health (ES026856, ES031529, GM070641, CA274603) and the National Research Foundation of Singapore through the Singapore-MIT Alliance for Research and Technology Antimicrobial Resistance Interdisciplinary Research Group. Quantitative graphics were generated in R and Morpheus, pictorial graphics were generated using Inkscape and Biorender. The graphical abstract and figure 7 were generated using BioRender.com.

## Supplementary materials

The Supplementary Material for this article can be found online at:

https://www.dropbox.com/scl/fo/cmzontjnfhucbpktmj7cg/AMLL9oLICbmpEaWTCtX32YY?rlkey=vbfxtfxf1sucjowy7vencnoq7&st=xluztvdn&dl=0

## TABLE LEGENDS

**Table 1.** Human tumor suppressors (TSGs) and oncogenes.

**Table 2.** Mouse tumor suppressors (TSGs) and oncogenes.

## SUPPLEMENTARY FIGURE LEGENDS

**Supplementary Figure S1. Human oncogenes have distinct codon usage patterns relative to human tumor suppressor genes.** Codon usage bias in human oncogenes and tumor suppressor genes was quantified using T-statistics, which capture deviations in codon usage relative to the genomic background. (A) Heat map showing ICF-based T-statistics, where each row represents the group of oncogenes or tumor suppressor gene and each column represents codon. Values reflect relative enrichment or depletion of specific codons in each gene list. Hierarchical clustering was performed on both oncogenes and TSG, revealing that they segregate into distinct clusters, indicating systematic differences in isoacceptor codon usage preferences. (B) Heat map showing TCF-based T-statistics, summarizing overall codon usage bias across all codons for each gene. Hierarchical clustering based on total T-statistics further separates oncogenes from tumor suppressor genes, demonstrating that oncogenes occupy a distinct codon usage space at the global level.

**Supplementary Figure S2. Reproducibility of oncogene- and tumor suppressor–specific codon usage patterns across independently curated gene sets.** Codon usage bias was reanalyzed using an independently curated, updated list of human oncogenes and tumor suppressor genes (2024) to assess the reproducibility of the observed codon usage patterns. Stacked bar plots showing codon trends between human oncogenes and TSG which were quantified using T-statistics and evaluated using both the isoacceptor codon frequency (ICF) and total codon frequency (TCF) metrics. Red bars represent human oncogenes, and green bars represent human TSGs. Codons are along the x-axis and t-statistics on y-axis, where positive and negative value identifies a group of genes that have values higher or lower than average values, respectively for a specific codon. The overlapping bars indicates the number of codons where the T-statistic is in same positive or negative direction for both oncogenes and TSGs, while opposing indicates codon usage in opposite directions. (A) ICF- and (B) TCF-based T-statistics. The strong concordance in codon usage trends across independently curated gene lists demonstrates the robustness and reproducibility of the oncogene- and tumor suppressor–specific codon usage signatures.

**Supplementary Figure S3. K-means clustering reveals two distinct codon usage classes among human oncogenes and TSGs.** K-means clustering was performed on codon usage features using Z-score values (ICF) of (A) human oncogenes and (B) human TSGs to identify intrinsic patterns of synonymous codon preference. The analysis separated oncogenes and TSGs into two distinct clusters based on their codon usage profiles. One cluster (blue) is characterized by preferential use of G/C-ending codons (GC3), whereas the second cluster (red) preferentially overuses A/U-ending codons (AU3). Genes are colored coded according to cluster assignment, and clustering was driven by differences in third-position nucleotide composition across synonymous codons. Ellipses represent the 95% confidence intervals (CI) around the cluster centroids (mean codon usage profiles), illustrating the dispersion and separation of the two codon usage groups. These results indicate that oncogenes and TSGs are not homogeneous with respect to codon usage bias but instead segregate into discrete codon usage subclasses, suggesting potential differences in translational regulation or sensitivity to cellular tRNA availability.

**Supplementary Figure S4. Mouse oncogenes exhibit distinct codon usage patterns relative to mouse tumor suppressor genes and show conservation with human codon usage signatures.** (A) Heat map showing ICF T-statistics for mouse oncogenes and tumor suppressor genes. Rows represent the group of oncogenes or tumor suppressor gene(TSGs), and each column represents codon Values indicate relative enrichment or depletion of codons within each group. Hierarchical clustering was applied to both genes and codons, revealing distinct clustering of oncogenes and tumor suppressor genes based on isoacceptor-level codon usage. (B) Heat map showing TCF T-statistics for mouse genes, summarizing overall codon usage bias across all codons for each gene. Hierarchical clustering based on total T-statistics further separates mouse oncogenes from tumor suppressor genes, indicating global differences in codon usage patterns. (C) Comparative analysis of codon usage signatures between mouse and human oncogenes and tumor suppressor genes. Corresponding T-statistic trends across species demonstrate partial conservation of codon usage biases, supporting the presence of evolutionarily conserved codon usage features associated with oncogenic and tumor suppressive gene classes.

**Supplementary Figure S5. Human gene Ontology (GO) groups, oncogenes and tumor suppressors stratify using PCA analysis of ICF- based group measures.** (A) Principal component analysis (PCA) lattice plot showing pairwise scatter plots of PC1 to PC4 derived from the standardized T-statistic values from AmiGO2 for gene ontology groups and ICF measures for oncogenes and tumor suppressors. Each panel displays the distribution of genes along two principal components. The percentage of variance explained by each principal component is indicated on the diagonal panels. For example, panels in the top row correspond to PC1 on the y-axis. The panel in column 2 of that row therefore shows PC1 vs PC2 (PC2 on the x-axis, PC1 on the y-axis). Similarly, the panel in row 3, column 1 shows PC3 vs PC1. (B) Stacked bar graphs showing the mean principal component scores for PC1–PC4 calculated from the same PCA model. Scores are grouped by gene category: Gene Ontology (GO), oncogenes (red), and tumor suppressor genes (green). The bars represent group-wise mean PCA scores, summarizing the relative contribution of each group along the major axes of variance identified by PCA.

**Supplementary Figure S6. Gene Ontology (GO) based gene annotation of human oncogene and tumor suppressors for TCF based measures.** Heatmap of (A) Total (TCF) T-statistic values for 447 GOs, human oncogenes and human tumor suppressor genes. They were hierarchically clustered, describing codon usage patterns of all categories. (B) A three-dimensional (3D) PCA score plot was generated using the first three principal components (PC1, PC2, and PC3) and visualized as a two-dimensional (2D) projection at a fixed viewing angle. Each dot represents GOs and there are 6 cluster founds. Human oncogenes and tumor suppressors were labelled as black dots. (C) A cluster map depicting AU3 and GC3 bias among 6 clusters of human GOs, oncogenes and TSGs (Tumor suppressors), values corresponding to T-statistic with positive and negative values indicating enrichment or depletion respectively. Bias was considered significant at ±95% CI.

**Supplementary Figure S7. Gene Ontology (GO) based PCA analysis of human oncogene and tumor suppressors for TCF based measures.** (A) Principal component analysis (PCA) lattice plot showing pairwise scatter plots of PC1 to PC4 derived from the standardized T-statistic values from AmiGO2 for gene ontology groups and TCF measures for oncogenes and tumor suppressors. Each panel displays the distribution of genes along two principal components. The percentage of variance explained by each principal component is indicated on the diagonal panels. For example, panels in the top row correspond to PC1 on the y-axis. The panel in column 2 of that row therefore shows PC1 vs PC2 (PC2 on the x-axis, PC1 on the y-axis). Similarly, the panel in row 3, column 1 shows PC3 vs PC1. (B) Stacked bar graphs showing the mean principal component scores for PC1–PC4 calculated from the same PCA model. Scores are grouped by gene category: Gene Ontology (GO), oncogenes (red), and tumor suppressor genes (green). The bars represent group-wise mean PCA scores, summarizing the relative contribution of each group along the major axes of variance identified by PCA.

**Supplementary Figure S8. Gene Ontology (GO) based gene annotation of mouse oncogene and tumor suppressors for ICF based measures.** Heatmaps of (A) Isoacceptor (ICF) T-statistic values for 462 GOs, mouse oncogenes and mouse tumor suppressor genes. They were hierarchically clustered, describing codon usage patterns of all categories. (B) Principal component analysis (PCA) lattice plot showing pairwise scatter plots of PC1 to PC4 derived from the standardized T-statistic values from AmiGO2 for gene ontology groups and ICF measures for oncogenes and tumor suppressors.. Each panel displays the distribution of genes along two principal components. The percentage of variance explained by each principal component is indicated on the diagonal panels. For example, panels in the top row correspond to PC1 on the y-axis. The panel in column 2 of that row therefore shows PC1 vs PC2 (PC2 on the x-axis, PC1 on the y-axis). (C) Stacked bar graphs showing the mean principal component scores for PC1–PC4 calculated from the same PCA model. Scores are grouped by gene category: Gene Ontology (GO), oncogenes (red), and tumor suppressor genes (green). The bars represent group-wise mean PCA scores, summarizing the relative contribution of each group along the major axes of variance identified by PCA. (D) A three-dimensional PCA score plot was generated using the first three principal components (PC1, PC2, and PC3) and visualized as a two-dimensional projection at a fixed viewing angle. Each dot represents GOs, there are 5 cluster found. Mouse oncogenes and tumor suppressors were labelled as black dots.

**Supplementary Figure S9. Gene Ontology (GO) based gene annotation of mouse oncogene and tumor suppressors for TCF based measures.** Heatmaps of (A) Total (TCF) T-statistic values for 462 GOs, mouse oncogenes and mouse tumor suppressor genes. They were hierarchically clustered, describing codon usage patterns of all categories. (B) Principal component analysis (PCA) lattice plot showing pairwise scatter plots of PC1 to PC4 derived from the standardized T-statistic values from AmiGO2 for gene ontology groups and TCF measures for oncogenes and tumor suppressors.. Each panel displays the distribution of genes along two principal components. The percentage of variance explained by each principal component is indicated on the diagonal panels. For example, panels in the top row correspond to PC1 on the y-axis. The panel in column 2 of that row therefore shows PC1 vs PC2 (PC2 on the x-axis, PC1 on the y-axis). (C) Stacked bar graphs showing the mean principal component scores for PC1–PC4 calculated from the same PCA model. Scores are grouped by gene category: Gene Ontology (GO), oncogenes (red), and tumor suppressor genes (green). The bars represent group-wise mean PCA scores, summarizing the relative contribution of each group along the major axes of variance identified by PCA. (D) A three-dimensional PCA score plot was generated using the first three principal components (PC1, PC2, and PC3) and visualized as a two-dimensional projection at a fixed viewing angle. Each dot represents GOs, there are 5 cluster found. Mouse oncogenes and tumor suppressors were labelled as black dots.

## SUPPLEMENTARY FILES

**Also available at this dropbox link:** https://www.dropbox.com/scl/fo/0x5dap5mx7gnjpsczgpij/AIPWrajJoH9f0pKtzYG6U2Y?rlkey=9uq3xk5xpf71ena2oeajxx9wb&st=nai97hp2&dl=0

**Supplementary Table S1A:** ICF data including t-statistics and p-values for 66 human oncogenes and 73 tumor suppressors.

**Supplementary Table S1B:** Recent gene list from 2024_ICF data including t-statistics and p-values for 54 human oncogenes and 71 tumor suppressors.

**Supplementary Table S2A:** TCF data including t-statistics and p-values for 66 human oncogenes and 73 tumor suppressors.

**Supplementary Table S2B:** Recent gene list from 2024_TCF data including t-statistics and p-values for 54 human oncogenes and 71 tumor suppressors.

**Supplementary Table S3**: ICF Z-score data for human oncogenes

**Supplementary Table S4:** TCF Z-score data for human oncogenes

**Supplementary Table S5:** Average frequencies and Zscore for human oncogenes and TSG (ICF and TCF)

**Supplementary Table S6A:** Gene-specific average Z-score of human oncogenes for isoacceptor codon frequency (ICF)

**Supplementary Table S6B:** Gene-specific average Z-score of human oncogenes for total codon frequency (TCF)

**Supplementary Table S6C:** Gene-specific average frequencies of human oncogenes for isoacceptor codon frequency (ICF)

**Supplementary Table S6D:** Gene-specific average frequencies of human oncogenes for total codon frequency (TCF)

**Supplementary Table S7A:** Gene-specific average Z-score of human TSGs for isoacceptor codon frequency (ICF)

**Supplementary Table S7B:** Gene-specific average Z-score of human TSGs for total codon frequency (TCF)

**Supplementary Table S7C:** Gene-specific average frequencies of human TSGs for isoacceptor codon frequency (ICF)

**Supplementary Table S7D:** Gene-specific average frequencies of human TSGs for total codon frequency (TCF)

**Supplementary Table S8:** ICF data including t-statistics and p-values for 45 mouse oncogenes and 55 tumor suppressors.

**Supplementary Table S9:** TCF data including t-statistics and p-values for 45 mouse oncogenes and 55 tumor suppressors.

**Supplementary Table S10:** ICF Z-score data of mouse oncogenes.

**Supplementary Table S11:** TCF Z-score data of mouse oncogenes.

**Supplementary Table S12:** T-statistics data (both ICF and TCF) of oncogenes and tumor suppressors for both human and mouse, hierarchically clustered together and visualized as a heat map.

**Supplementary Table S13:** T-statistics of all human GOs for both ICF and TCF from AmiGO2 browser.

**Supplementary Table S14:** T-statistics of all mouse GOs for both ICF and TCF from AmiGO2 browser.

---

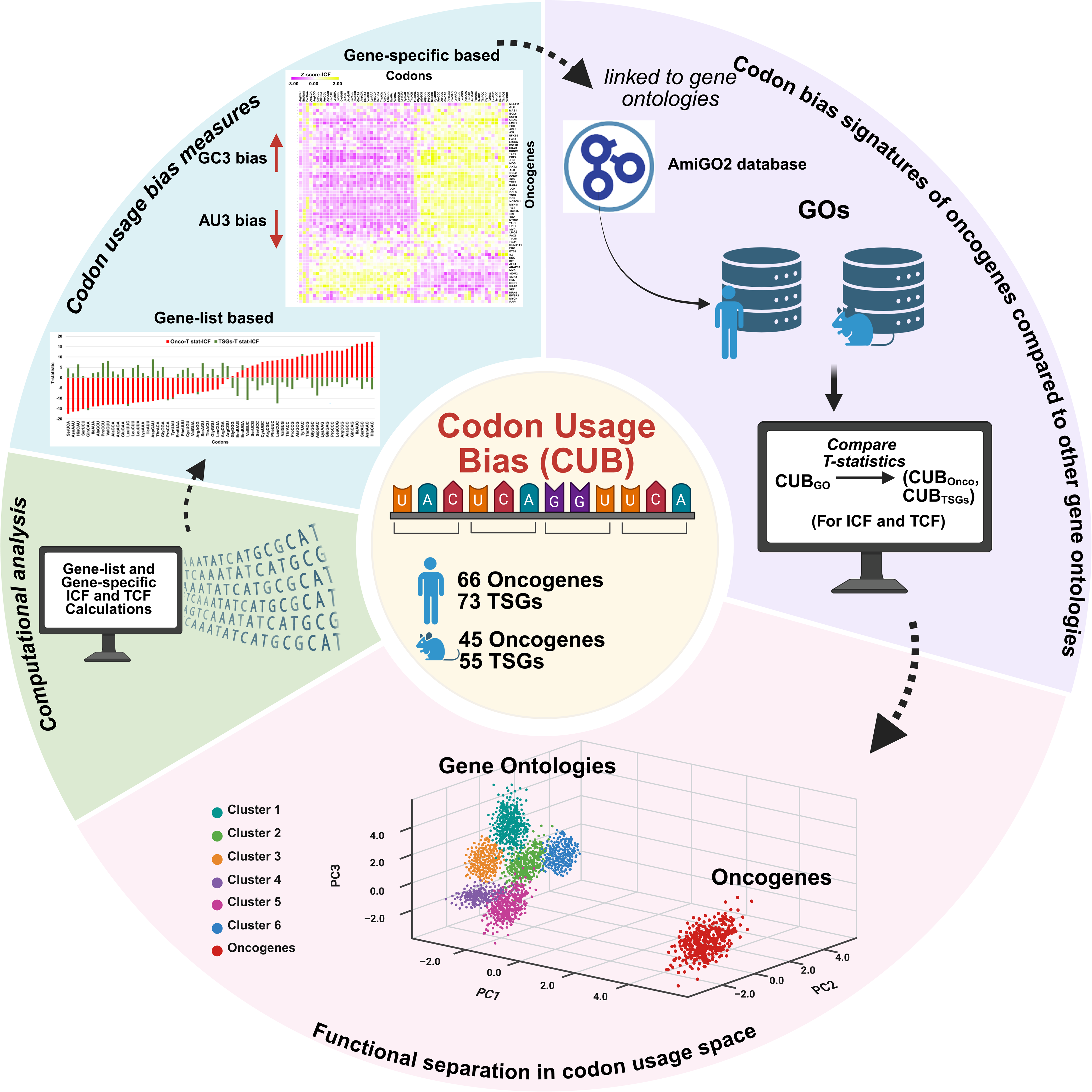

## REFERENCES

Agris, P.F. (2004). Decoding the genome: a modified view. Nucleic acids research 32(1), 223–238. doi: 10.1093/nar/gkh185.

Agris, P.F., Vendeix, F.A., Graham, W.D. (2007). tRNA’s wobble decoding of the genome: 40 years of modification. Journal of molecular biology 366(1), 1–13. doi: 10.1016/j.jmb.2006.11.046.

Aleksander, S.A., Balhoff, J., Carbon, S., Cherry, J.M., Drabkin, H.J., Ebert, D., Feuermann, M., Gaudet, P., Harris, N.L., Hill, D.P. (2023). The gene ontology knowledgebase in 2023. Genetics 224(1), iyad031. doi: 10.1093/genetics/iyad031.

Añazco-Guenkova, A.M., Miguel-López, B., Monteagudo-García, Ó., García-Vílchez, R., Blanco, S. (2024). The impact of tRNA modifications on translation in cancer: identifying novel therapeutic avenues. NAR cancer 6(1), zcae012. doi: 10.1093/narcan/zcae012.

Ashburner, M., Ball, C.A., Blake, J.A., Botstein, D., Butler, H., Cherry, J.M., Davis, A.P., Dolinski, K., Dwight, S.S., Eppig, J.T., Harris, M.A. (2000). Gene ontology: tool for the unification of biology. Nature genetics 25(1), 25–29. doi: 10.1038/75556.

Ashraf, S.S., Ansari, G., Guenther, R., Sochacka, E., Malkiewicz, A., Agris, P.F. (1999). The uridine in "U-turn": contributions to tRNA-ribosomal binding. RNA 5(4), 503–511. doi: 10.1017/s1355838299981931.

Ashraf, S.S., Guenther, R.H., Ansari, G., Malkiewicz, A., Sochacka, E., Agris, P.F. (2000). Role of modified nucleosides of yeast tRNA(Phe) in ribosomal binding. Cell Biochemistry and Biophysics 33(3), 241–252. doi: 10.1385/CBB:33:3:241.

Barbieri, I., Kouzarides, T. (2020). Role of RNA modifications in cancer. Nature Reviews Cancer 20(6), 303–322. doi: 10.1038/s41568-020-0253-2.

Barna, M., Pusic, A., Zollo, O., Costa, M., Kondrashov, N., Rego, E., Rao, P.H., Ruggero, D. (2008). Suppression of Myc oncogenic activity by ribosomal protein haploinsufficiency. Nature 456(7224), 971–975. doi: 10.1038/nature07449.

Barrington, C.L., Galindo, G., Koch, A.L., Horton, E.R., Morrison, E.J., Tisa, S., Stasevich, T.J., Rissland, O.S. (2023). Synonymous codon usage regulates translation initiation. Cell reports 42(12). doi: 10.1016/j.celrep.2023.113413.

Begley, T.J., Rosenbach, A.S., Ideker, T., Samson, L.D. (2002). Damage recovery pathways in Saccharomyces cerevisiae revealed by genomic phenotyping and interactome mapping. Molecular cancer research 1(2), 103–112.

Begley, U., Dyavaiah, M., Patil, A., Rooney, J.P., DiRenzo, D., Young, C.M., Conklin, D.S., Zitomer, R.S., Begley, T.J. (2007). Trm9-catalyzed tRNA modifications link translation to the DNA damage response. Molecular cell 28(5), 860–870. doi: 10.1016/j.molcel.2007.09.021

Begley, U., Sosa, M.S., Avivar-Valderas, A., Patil, A., Endres, L., Estrada, Y., Chan, C.T., Su, D., Dedon, P.C., Aguirre-Ghiso, J.A., Begley, T. (2013). A human tRNA methyltransferase 9-like protein prevents tumour growth by regulating LIN9 and HIF1-alpha. EMBO molecular medicine 5(3), 366–368. doi: 10.1002/emmm.201201161.

Beroukhim, R., Mermel, C.H., Porter, D., Wei, G., Raychaudhuri, S., Donovan, J., Barretina, J., Boehm, J.S., Dobson, J., Urashima, M., Mc Henry, K.T. (2010). The landscape of somatic copy-number alteration across human cancers. Nature 463(7283), 899–905. doi: 10.1038/nature08822.

Boccaletto, P., Stefaniak, F., Ray, A., Cappannini, A., Mukherjee, S., Purta, E., Kurkowska, M., Shirvanizadeh, N., Destefanis, E., Groza, P., Avşar, G. (2022). MODOMICS: a database of RNA modification pathways. 2021 update. Nucleic acids research 50(D1), D231-D235. doi: 10.1093/nar/gkab1083.

Carbon, S., Ireland, A., Mungall, C.J., Shu, S., Marshall, B., Lewis, S., AmiGO Hub, Web Presence Working Group (2009). AmiGO: online access to ontology and annotation data. Bioinformatics 25(2), 288–289. doi: 10.1093/bioinformatics/btn615.

Carroll, M., Borden, K.L. (2013). The oncogene eIF4E: using biochemical insights to target cancer. Journal of Interferon & Cytokine Research 33(5), 227–238. doi: 10.1089/jir.2012.0142.

Chan, C., Pham, P., Dedon, P.C., Begley, T.J. (2018). Lifestyle modifications: coordinating the tRNA epitranscriptome with codon bias to adapt translation during stress responses. Genome biology 19(1), 228. doi: 10.1186/s13059-018-1611-1.

Chan, C.T., Dyavaiah, M., DeMott, M.S., Taghizadeh, K., Dedon, P.C., Begley, T.J. (2010). A quantitative systems approach reveals dynamic control of tRNA modifications during cellular stress. PLoS genetics 6(12), e1001247. doi: 10.1371/journal.pgen.1001247.

Chan, C.T., Pang, Y.L.J., Deng, W., Babu, I.R., Dyavaiah, M., Begley, T.J., Dedon, P.C. (2012). Reprogramming of tRNA modifications controls the oxidative stress response by codon-biased translation of proteins. Nat. Commun. 3(1), 937. doi: 10.1038/ncomms1938.

Chionh, Y.H., McBee, M., Babu, I.R., Hia, F., Lin, W., Zhao, W., Cao, J., Dziergowska, A., Malkiewicz, A., Begley, T.J., Alonso, S., Dedon, P.C. (2016). tRNA-mediated codon-biased translation in mycobacterial hypoxic persistence. Nature communications 7(1), 13302. doi: 10.1038/ncomms13302.

Consortium, T.U. (2025). UniProt: the Universal Protein Knowledgebase in 2025. Nucleic Acids Research volume 53(D1), D609–D617. doi: 10.1093/nar/gkae1010.

Cui, L., Ma, R., Cai, J., Guo, C., Chen, Z., Yao, L., Wang, Y., Fan, R., Wang, X., Shi, Y. (2022). RNA modifications: importance in immune cell biology and related diseases. Signal transduction and targeted therapy 7(1), 334. doi: 10.1038/s41392-022-01175-9.

Dai, Z., Liu, H., Liao, J., Huang, C., Ren, X., Zhu, W., Zhu, S., Peng, B., Li, S., Lai, J., Liang, L. (2021). N7- Methylguanosine tRNA modification enhances oncogenic mRNA translation and promotes intrahepatic cholangiocarcinoma progression. Molecular cell 81(16), 3339–3355. doi: 10.1016/j.molcel.2021.07.003

Dang, C.V. (2012). MYC on the path to cancer. Cell 149(1), 22–35. doi: 10.1016/j.cell.2012.03.003.

Davis, E.T., Raman, R., Byrne, S.R., Ghanegolmohammadi, F., Mathur, C., Begley, U., Dedon, P.C., Begley, T.J. (2025). Genes and Pathways Comprising the Human and Mouse ORFeomes Display Distinct Codon Bias Signatures that Can Regulate Protein Levels. bioRxiv. doi: 10.1101/2025.02.03.636209.

De Jong, M., Maina, T. (2010). Of mice and humans: are they the same?—Implications in cancer translational research. Journal of Nuclear Medicine 51(4), 501–504. doi: 10.2967/jnumed.109.065706.

Dedon, P.C., Begley, T.J. (2014). A system of RNA modifications and biased codon use controls cellular stress response at the level of translation. Chemical research in toxicology 27(3), 330–337. doi: 10.1021/tx400438d.

Dedon, P.C., Begley, T.J. (2022). Dysfunctional tRNA reprogramming and codon-biased translation in cancer. Trends in molecular medicine 28(11), 964–978. doi: 10.1016/j.molmed.2022.09.007.

Delaunay, S., Rapino, F., Tharun, L., Zhou, Z., Heukamp, L., Termathe, M., Shostak, K., Klevernic, I., Florin, A., Desmecht, H., Desmet, C.J. (2016). Elp3 links tRNA modification to IRES-dependent translation of LEF1 to sustain metastasis in breast cancer. Journal of Experimental Medicine 213(11), 2503–2523. doi: 10.1084/jem.20160397.

Dempster, A.P., Laird, N.M., Rubin, D.B. (1977). Maximum likelihood from incomplete data via the EM algorithm. Journal of the royal statistical society: series B (methodological) 39(1), 1–22. doi: 10.1111/j.2517-6161.1977.tb01600.x.

Dhanasekaran, R., Deutzmann, A., Mahauad-Fernandez, W.D., Hansen, A.S., Gouw, A.M., Felsher, D.W. (2022). The MYC oncogene - the grand orchestrator of cancer growth and immune evasion. Nature reviews Clinical oncology 19(1), 23–26. doi: 10.1038/s41571-021-00549-2.

Dick, F.A., Rubin, S.M. (2013). Molecular mechanisms underlying RB protein function Nature reviews Molecular cell biology 14(5), 297–306. doi: 10.1038/nrm3567.

El Yacoubi, B., Bailly, M., de Crécy-Lagard, V. (2012). Biosynthesis and function of posttranscriptional modifications of transfer RNAs Annual review of genetics 46(1), 69–95. doi: 10.1146/annurev-genet-110711-155641.

Ellis, L.M., Fidler, I.J. (2010). Finding the tumor copycat: therapy fails, patients don’t. Nature Medicine 16(9), 974–975. doi: 10.1038/nm0910-974.

Endres, L., Begley, U., Clark, R., Gu, C., Dziergowska, A., Małkiewicz, A., Melendez, J.A., Dedon, P.C., Begley, T.J. (2015a). Alkbh8 regulates selenocysteine-protein expression to protect against reactive oxygen species damage. PloS one 10(7), e0131335. doi: 10.1371/journal.pone.0131335.

Endres, L., Dedon, P.C., Begley, T.J. (2015b). Codon-biased translation can be regulated by wobble-base tRNA modification systems during cellular stress responses. RNA biology 12(6), 603–614. doi: 10.1080/15476286.2015.1031947.

Endres, L., Fasullo, M., Rose, R. (2019). tRNA modification and cancer: potential for therapeutic prevention and intervention. Future medicinal chemistry 11(8), 885–900. doi: 10.4155/fmc-2018-0404.

Fraley, C., Raftery, A.E. (2002). Model-based clustering, discriminant analysis, and density estimation. Journal of the American statistical Association 97(458), 611–631. doi: 10.1198/016214502760047131.

Fu, J., Dang, Y., Counter, C., Liu, Y. (2018). Codon usage regulates human KRAS expression at both transcriptional and translational levels. Journal of Biological Chemistry 293(46), 17929–17940. doi: 10.1074/jbc.RA118.004908.

Ghanegolmohammadi, F., Ohnuki, S., Byrne, S., Raman, R., Begley, T.J., Dedon, P.C. (2026). Synonymous codon usage defines functional gene families. BMC biology 24(1), 44. doi: 10.1186/s12915-026-02505-x.

Gingold, H., Tehler, D., Christoffersen, N.R., Nielsen, M.M., Asmar, F., Kooistra, S.M., Christophersen, N.S., Christensen, L.L., Borre, M., Sørensen, K.D., Andersen, L.D. (2014). A dual program for translation regulation in cellular proliferation and differentiation. Cell 158(6), 1281–1292. doi: 10.1016/j.cell.2014.08.011.

Goodarzi, H., Nguyen, H.C., Zhang, S., Dill, B.D., Molina, H., Tavazoie, S.F. (2016). Modulated expression of specific tRNAs drives gene expression and cancer progression. Cell 165(6), 1416–1427. doi: 10.1016/j.cell.2016.05.046

Gu, C., Begley, T.J., Dedon, P.C. (2014). tRNA modifications regulate translation during cellular stress. FEBS letters 588(23), 4287–4296. doi: 10.1016/j.febslet.2014.09.038.

Han, H., Yang, C., Ma, J., Zhang, S., Zheng, S., Ling, R., Sun, K., Guo, S., Huang, B., Liang, Y., Wang, L. (2022). N7-methylguanosine tRNA modification promotes esophageal squamous cell carcinoma tumorigenesis via the RPTOR/ULK1/autophagy axis. Nature communications 13(1), 1478. doi: 10.1038/s41467-022-29125-7.

Hanahan, D., Weinberg, R.A. (2011). Hallmarks of cancer: the next generation. Cell 144(5), 646–674. doi: 10.1016/j.cell.2011.02.013

Hanson, G., Coller, J. (2018). Codon optimality, bias and usage in translation and mRNA decay. Nature reviews Molecular cell biology 19, 20–30. doi: 10.1038/nrm.2017.91.

Hayden, E.C. (2008). Cancer complexity slows quest for cure. Nature 455(7210), 148–149.

Hershey, J.W., Sonenberg, N., Mathews, M.B. (2012). Principles of translational control: an overview. Cold Spring Harbor perspectives in biology 4(12), a011528. doi: 10.1101/cshperspect.a011528.

Hsieh, A.C., Liu, Y., Edlind, M.P., Ingolia, N.T., Janes, M.R., Sher, A., Shi, E.Y., Stumpf, C.R., Christensen, C., Bonham, M.J., Wang, S. (2012). The translational landscape of mTOR signalling steers cancer initiation and metastasis. . Nature 485(7396), 55-61. doi: 10.1038/nature10912.

Huber, S.M., Begley, U., Sarkar, A., Gasperi, W., Davis, E.T., Surampudi, V., Lee, M., Melendez, J.A., Dedon, P.C., Begley, T.J. (2022). Arsenite toxicity is regulated by queuine availability and oxidation-induced reprogramming of the human tRNA epitranscriptome. Proceedings of the National Academy of Sciences 119(38), e2123529119. doi: 10.1073/pnas.2123529119.

Jolliffe, I. (2025). "Principal Component Analysis " in International Encyclopedia of Statistical Science. (Berlin, Heidelberg: Springer), 1945-1948.

Jolliffe, I.T., Cadima, J. (2016). Principal component analysis: a review and recent developments. *Philosophical transactions of the royal society A: Mathematical*, Physical and Engineering Sciences 374(2065), 20150202. doi: 10.1098/rsta.2015.0202.

Jonkhout, N., Tran, J., Smith, M.A., Schonrock, N., Mattick, J.S., Novoa, E.M. (2017). The RNA modification landscape in human disease. RNA 23(12), 1754–1769. doi: 10.1261/rna.063503.117.

Jungfleisch, J., Böttcher, R., Talló-Parra, M., Pérez-Vilaró, G., Merits, A., Novoa, E.M., Díez, J. (2022). CHIKV infection reprograms codon optimality to favor viral RNA translation by altering the tRNA epitranscriptome. Nature communications, 13(1), 4725. doi: 10.1038/s41467-022-31835-x.

Kissil, J.L., Walmsley, M.J., Hanlon, L., Haigis, K.M., Bender Kim, C.F., Sweet-Cordero, A., Eckman, M.S., Tuveson, D.A., Capobianco, A.J., Tybulewicz, V.L., Jacks, T. (2007). Requirement for Rac1 in a K-ras–induced lung cancer in the mouse. Cancer research 67(17), 8089–8094. doi: 10.1158/0008-5472.CAN-07-2300.

Kovalski, J.R., Kuzuoglu-Ozturk, D., Ruggero, D. (2022). Protein synthesis control in cancer: selectivity and therapeutic targeting. The EMBO Journal 41(8), EMBJ2021109823. doi: 10.15252/embj.2021109823.

Ladang, A., Rapino, F., Heukamp, L.C., Tharun, L., Shostak, K., Hermand, D., Delaunay, S., Klevernic, I., Jiang, Z., Jacques, N., Jamart, D. (2015). Elp3 drives Wnt-dependent tumor initiation and regeneration in the intestine. Journal of Experimental Medicine 212(12), 2057–2075.

Lampson, B.L., Pershing, N.L., Prinz, J.A., Lacsina, J.R., Marzluff, W.F., Nicchitta, C.V., MacAlpine, D.M., Counter, C.M. (2013). Rare codons regulate KRas oncogenesis Current Biology 23(1), 70–75. doi: 10.1016/j.cub.2012.11.031.

Leonardi, A., Evke, S., Lee, M., Melendez, J.A., Begley, T.J. (2019). Epitranscriptomic systems regulate the translation of reactive oxygen species detoxifying and disease linked selenoproteins. Free Radical Biology and Medicine 143, 573–593. doi: 10.1016/j.freeradbiomed.2019.08.030.

Levine, A.J., Oren, M. (2009). The first 30 years of p53: growing ever more complex. Nature reviews cancer 9(10), 749–758. doi: 10.1038/nrc2723.

Levine, D.A., Cancer genome atlas research network. (2013). Integrated genomic characterization of endometrial carcinoma. Nature 497, 67-73. doi: 10.1038/nature12113.

Li, J., Wang, L., Hahn, Q., Nowak, R.P., Viennet, T., Orellana, E.A., Roy Burman, S.S., Yue, H., Hunkeler, M., Fontana, P., Wu, H. (2023). Structural basis of regulated m7G tRNA modification by METTL1–WDR4. Nature 613(7943), 391–397. doi: 10.1038/s41586-022-05566-4.

Li, Q., Li, J., Yu, C.P., Chang, S., Xie, L.L., Wang, S. (2021). Synonymous mutations that regulate translation speed might play a non-negligible role in liver cancer development. BMC cancer 21(1), 388. doi: 10.1186/s12885-021-08131-w.

Lin, S., Liu, Q., Lelyveld, V.S., Choe, J., Szostak, J.W., Gregory, R.I (2018). Mettl1/Wdr4-mediated m7G tRNA methylome is required for normal mRNA translation and embryonic stem cell self-renewal and differentiation. Molecular cell 71(2), 244–255. doi: 10.1016/j.molcel.2018.06.001

Meyer, K.D., Jaffrey, S.R. (2017). Rethinking m6A readers, writers, and erasers. Annual review of cell and developmental biology 33, 319–342. doi: 10.1146/annurev-cellbio-100616-060758.

Mitchener, M.M., Begley, T.J., Dedon, P.C. (2023). Molecular coping mechanisms: reprogramming tRNAs to regulate codon-biased translation of stress response proteins. Accounts of Chemical Research 56(23), 3504–3514. doi: 10.1021/acs.accounts.3c00572.

Network, C.G.A. (2012). Comprehensive molecular portraits of human breast tumors. Nature 490(7418), 61–70. doi: 10.1038/nature11412.

Orellana, E.A., Liu, Q., Yankova, E., Pirouz, M., De Braekeleer, E., Zhang, W., Lim, J., Aspris, D., Sendinc, E., Garyfallos, D.A., Gu, M. (2021). METTL1-mediated m7G modification of Arg-TCT tRNA drives oncogenic transformation. Molecular cell 81(16), 3323–3338. doi: 10.1016/j.molcel.2021.06.031.

Pavon-Eternod, M., Gomes, S., Geslain, R., Dai, Q., Rosner, M.R., Pan, T. (2009). tRNA over-expression in breast cancer and functional consequences. Nucleic acids research 37(21), 7268–7280. doi: 10.1093/nar/gkp787.

Pedersen, T. (2024). "ggforce: Accelerating “ggplot2”[R Package]". 0.4.2. ed.: Comprehensive R Archive Network (CRAN)).

Pershing, N.L., Lampson, B.L., Belsky, J.A., Kaltenbrun, E., MacAlpine, D.M., Counter, C.M. (2015). Rare codons capacitate Kras-driven de novo tumorigenesis The Journal of clinical investigation 125(1), 222–233. doi: 10.1172/JCI77627.

Plotkin, J.B., Kudla, G. (2011). Synonymous but not the same: The causes and consequences of codon bias. Nature Reviews Genetics 12(1), 32–42. doi: 10.1038/nrg2899.

Presnyak, V., Alhusaini, N., Chen, Y.H., Martin, S., Morris, N., Kline, N., Olson, S., Weinberg, D., Baker, K.E., Graveley, B.R., Coller, J. (2015). Codon optimality is a major determinant of mRNA stability Cell 160(6), 1111–1124. doi: 10.1016/j.cell.2015.02.029.

Pylayeva-Gupta, Y., Grabocka, E., Bar-Sagi, D. (2011). RAS oncogenes: weaving a tumorigenic web. Nature Reviews Cancer 11(11), 761–774. doi: 10.1038/nrc3106.

Rangarajan, A., Weinberg, R.A. (2003). Comparative biology of mouse versus human cells: modelling human cancer in mice. Nature Reviews Cancer 3(12), 952–959. doi: 10.1038/nrc1235.

Rapino, F., Delaunay, S., Rambow, F., Zhou, Z., Tharun, L., De Tullio, P., Sin, O., Shostak, K., Schmitz, S., Piepers, J., Ghesquière, B. (2018). Codon specific translation reprogramming promotes resistance to targeted therapy. Nature 558(7711), 605–609. doi: 10.1038/s41586-018-0243-7.

Rapino, F., Delaunay, S., Zhou, Z., Chariot, A., Close, P. (2017). tRNA Modification: Is Cancer Having a Wobble? Trends in cancer 3(4), 249-252. doi: 10.1016/j.trecan.2017.02.004

Robichaud, N., Sonenberg, N., Ruggero, D., Schneider, R.J. (2019). Translational control in cancer. Cold Spring Harbor perspectives in biology 11(7), a032896. doi: 10.1101/cshperspect.a032896.

Roundtree, I.A., Evans, M.E., Pan, T., He, C. (2017). Dynamic RNA modifications in gene expression regulation. Cell 169(7), 1187–1200. doi: 10.1016/j.cell.2017.05.045.

Ruggero, D. (2009). The role of Myc-induced protein synthesis in cancer Cancer research 69(23), 8839–8843. doi: 10.1158/0008-5472.CAN-09-1970.

Saci, A., Cantley, L.C., Carpenter, C.L. (2011). Rac1 regulates the activity of mTORC1 and mTORC2 and controls cellular size. Molecular cell 42(1), 50–61. doi: 10.1016/j.molcel.2011.03.017

Sajwan, R., Wang, L., Casar-Borota, O., Karakostis, K., Chen, S., Fahraeus, R., Gu, X., Vadivel Gnanasundram, S. (2025). A cancer-associated TP53 synonymous mutation induces synthesis of the p53 isoform p53/47: Cellular and Molecular Biology. British Journal of Cancer 133(7), 970–975. doi: 10.1038/s41416-025-03127-w.

Seres, M., Spacayova, K., Sulova, Z., Spaldova, J., Breier, A., Pavlikova, L. (2025). Dynamic multilevel regulation of EGFR, KRAS, and MYC oncogenes: driving cancer cell proliferation through (Epi) genetic and post-transcriptional/translational pathways. . Cancers 17(2), 248. doi: 10.3390/cancers17020248.

Shan, K.S., Rehman, T.U., Ivanov, S., Domingo, G., Raez, L.E. (2024). Molecular targeting of the BRAF proto-oncogene/mitogen-activated protein kinase (MAPK) pathway across cancers. . International Journal of Molecular Sciences 25(1), 624. doi: 10.3390/ijms25010624.

Sharp, P.M., Li, W.H. (1987). The codon adaptation index—a measure of directional synonymous codon usage bias, and its potential applications. Nucleic acids research 15(3), 1281–1295. doi: 10.1093/nar/15.3.1281.

Shi, H., Wei, J., He, C. (2019). Where, when, and how: context-dependent functions of RNA methylation writers, readers, and erasers Molecular cell 74(4), 640–650. doi: 10.1016/j.molcel.2019.04.025.

Song, M.S., Salmena, L., Pandolfi, P.P. (2012). The functions and regulation of the PTEN tumour suppressor. Nature reviews Molecular cell biology 13(5), 283–296. doi: 10.1038/nrm3330.

Stojchevski, R., Sutanto, E.A., Sutanto, R., Hadzi-Petrushev, N., Mladenov, M., Singh, S.R., Sinha, J.K., Ghosh, S., Yarlagadda, B., Singh, K.K., Verma, P. (2025). Translational advances in oncogene and tumor-suppressor gene research. Cancers 17(6), 1008. doi: 10.3390/cancers17061008.

Supek, F., Miñana, B., Valcárcel, J., Gabaldón, T., Lehner, B. (2014). Synonymous mutations frequently act as driver mutations in human cancers. Cell 156(6), 1324–1335. doi: 10.1016/j.cell.2014.01.051

Suzuki, T. (2021). The expanding world of tRNA modifications and their disease relevance. Nature Reviews Molecular Cell Biology 22(6), 375–392. doi: 10.1038/s41580-021-00342-0.

Thoreen, C.C., Chantranupong, L., Keys, H.R., Wang, T., Gray, N.S., Sabatini, D.M. (2012). A unifying model for mTORC1-mediated regulation of mRNA translation. Nature 485(7396), 109–113. doi: 10.1038/nature11083.

Thoreen, C.C., Kang, S.A., Chang, J.W., Liu, Q., Zhang, J., Gao, Y., Reichling, L.J., Sim, T., Sabatini, D.M., Gray, N.S. (2009). An ATP-competitive mammalian target of rapamycin inhibitor reveals rapamycin-resistant functions of mTORC1. Journal of Biological Chemistry 284(12), 8023–8032. doi: 10.1074/jbc.M900301200.

Torres, A.G., Batlle, E., de Pouplana, L.R. (2014). Role of tRNA modifications in human diseases. Trends in Molecular Medicine 20(6), 306–314. doi: 10.1016/j.molmed.2014.01.008

Trexler, M., Bányai, L., Kerekes, K., Patthy, L. (2024). Arginines of the CGN codon family are Achilles’ heels of cancer genes. Scientific Reports 14(1), 11715. doi: 10.1038/s41598-024-62553-7.

Van Riggelen, J., Yetil, A., Felsher, D.W. (2010). MYC as a regulator of ribosome biogenesis and protein synthesis. Nature Reviews Cancer 10(4), 301–309. doi: 10.1038/nrc2819.

Väre, V.Y., Eruysal, E.R., Narendran, A., Sarachan, K.L., Agris, P.F. (2017). Chemical and conformational diversity of modified nucleosides affects tRNA structure and function. Biomolecules 7(1), 29. doi: 10.3390/biom7010029.

Varghese, A.M., Perry, M.A., Chou, J.F., Nandakumar, S., Muldoon, D., Erakky, A., Zucker, A., Fong, C., Mehine, M., Nguyen, B., Basturk, O. (2025). Clinicogenomic landscape of pancreatic adenocarcinoma identifies KRAS mutant dosage as prognostic of overall survival. Nature medicine 31(2), 466–477. doi: 10.1038/s41591-024-03362-3.

Walker, E.J., Zhang, C., Castelo-Branco, P., Hawkins, C., Wilson, W., Zhukova, N., Alon, N., Novokmet, A., Baskin, B., Ray, P., Knobbe, C. (2012). Monoallelic expression determines oncogenic progression and outcome in benign and malignant brain tumors. Cancer research 72(3), 636–644. doi: 10.1158/0008-5472.CAN-11-2266.

Wang, X., Lu, Z., Gomez, A., Hon, G.C., Yue, Y., Han, D., Fu, Y., Parisien, M., Dai, Q., Jia, G., Ren, B. (2014). N 6-methyladenosine-dependent regulation of messenger RNA stability. Nature 505(7481), 117–120. doi: 10.1038/nature12730.

Wang, X., Zhao, B.S., Roundtree, I.A., Lu, Z., Han, D., Ma, H., Weng, X., Chen, K., Shi, H., He, C. (2015). N6-methyladenosine modulates messenger RNA translation efficiency Cell 161(6), 1388-1399. doi: 10.1016/j.cell.2015.05.014.

Wu, X., Xu, M., Yang, J.R., Lu, J. (2024). Genome-wide impact of codon usage bias on translation optimization in Drosophila melanogaster. Nature communications 15(1), 8329. doi: 10.1038/s41467-024-52660-4.

Xiao, W., Adhikari, S., Dahal, U., Chen, Y.S., Hao, Y.J., Sun, B.F., Sun, H.Y., Li, A., Ping, X.L., Lai, W.Y., Wang, X. (2016). Nuclear m6A reader YTHDC1 regulates mRNA splicing. Molecular cell 61(4), 507–519. doi: 10.1016/j.molcel.2016.01.012

Yang, X., Wu, H. (2024). RAS signaling in carcinogenesis, cancer therapy and resistance mechanisms. Journal of Hematology & Oncology 17(1), 108. doi: 10.1186/s13045-024-01631-9.

Zhang, C., Samanta, D., Lu, H., Bullen, J.W., Zhang, H., Chen, I., He, X., Semenza, G.L. (2016). Hypoxia induces the breast cancer stem cell phenotype by HIF-dependent and ALKBH5-mediated m6A-demethylation of NANOG mRNA. Proceedings of the National Academy of Sciences 113(14), E2047–E2056. doi: 10.1073/pnas.1602883113.

Zhang, W., Foo, M., Eren, A.M., Pan, T. (2022). tRNA modification dynamics from individual organisms to metaepitranscriptomics of microbiomes. Molecular Cell 82(5), 891–906. doi: 10.1016/j.molcel.2021.12.007

Zhou, Y., Zhou, B., Pache, L., Chang, M., Khodabakhshi, A.H., Tanaseichuk, O., Benner, C., Chanda, S.K. (2019). Metascape provides a biologist-oriented resource for the analysis of systems-level datasets. Nature communications 10(1), 1523. doi: 10.1038/s41467-019-09234-6.

