## Supplementary Table 1 and 2 for "Oncogenes have the most distinct codon biases in the genome and codon signatures that oppose tumor suppressor genes"

Table 1: Human tumor suppressors (TSGs) and oncogenes.

| **TUMOR SUPPRESSORS (TSGs)** | | | | | **ONCOGENES** | | | | |
|---|---|---|---|---|---|---|---|---|---|
| APC | CDKN2C | IL2 | NOTCH1 | SMARCB1 | ABL1 | ERG | LMO1 | NRAS | TAL1 |
| ARHGEF12 | CEBPA | JAK1 | NPM1 | SOCS1 | AFF4 | ETS1 | LMO2 | NTRK1 | TAL2 |
| ATF3 | CREB1 | JAK2 | NR4A3 | STK11 | AKAP13 | EWSR1 | LYL1 | PAX5 | TCF3 |
| ATM | CREBBP | JAK3 | NUP98 | SUFU | AKT2 | FES | MAS1 | PBX1 | TIAM1 |
| BCL11B | CYLD | JAKMIP1 | PALB2 | SYK | ALK | FGF3 | MCF2 | PIM1 | TLX1 |
| BMPR1A | DDX5 | JAKMIP2 | PML | TCF3 | AXL | FGF4 | MCF2L | RAF1 | TSC2 |
| BRCA1 | EPB41L3 | JAKMIP3 | PMS1 | TNFAIP3 | BCL2 | FOS | MDM2 | RARA | |
| BRCA2 | EXT1 | KLKB1 | PMS2 | TP53 | BCL3 | GLI1 | MLLT11 | REL | |
| CARS1 | EXT2 | MAP2K4 | PTEN | TSC1 | BCL6 | GNAS | MOS | RET | |
| CARS2 | FBXW7 | MDM4 | RB1 | TSC2 | BCR | HRAS | MYB | ROS1 | |
| CBFA2T3 | FH | MEN1 | RUNX1 | VHL | CCND1 | IL3 | MYCL | RUNX1 | |
| CDH1 | FLT3 | MLH1 | SDHB | WRN | CSF1R | JUN | MYCN | RUNX1T1 | |
| CDH11 | FOXP1 | MSH2 | SDHD | | DEK | KIT | MYH11 | SET | |
| CDKN1A | GPC3 | NF1 | SMAD2 | | EGFR | KRAS | NFKB2 | SKI | |
| CDKN2A | IDH1 | NF2 | SMARCA4 | | ERBB2 | LCK | NOTCH1 | SRC | |

Table 2: Mouse tumor suppressors (TSGs) and oncogenes.

| TUMOR SUPPRESSORS (TSGs) | | | | | ONCOGENES | | | |
|---|---|---|---|---|---|---|---|---|
| APC | CHD5 | MAFB | RASSF2 | TSC2 | ABL1 | FLT3 | MOS | SRC |
| ATM | DLEC1 | MAPKAPK5 | RASSF4 | VHL | ARAF | FYN | MYB | STYK1 |
| BCL10 | EPB41L3 | MCTS1 | RASSF5 | WT1 | AURKA | HCK | NOTCH4 | USP4 |
| BRCA1 | FES | NBL1 | RB1 | | BRAF | JUN | PDGFB | WNT1 |
| BRCA2 | FRK | NDRG2 | RB1CC1 | | CDK4 | KDSR | PDGFRA | WNT3 |
| BRMS1 | GPR68 | PARK7 | RBL1 | | CDT1 | KIT | PIM1 | YES1 |
| CADM1 | HIF3A | PDCD4 | RBL2 | | CSF1R | KRAS | PIM2 | |
| CADM4 | ING1 | PLK2 | SIK1 | | ETS1 | LCK | PIM3 | |
| CCAR2 | ING4 | PML | SIRT6 | | FER | LYN | PRKCI | |
| CDKN1A | LACTB | PMS2 | STK11 | | FES | MAP3K8 | PTP4A3 | |
| CDKN2A | LATS1 | PTEN | TRP53 | | FGF6 | MAS1 | RAF1 | |
| CDKN2B | LATS2 | RAP1A | TRP73 | | FGFR2 | MERTK | RET | |
| CDKN2D | MAF | RASSF1 | TSC1 | | FGR | MET | ROS1 | |
