## Supplementary figures and images for "Oncogenes have the most distinct codon biases in the genome and codon signatures that oppose tumor suppressor genes"

### Supplementary Figure 1

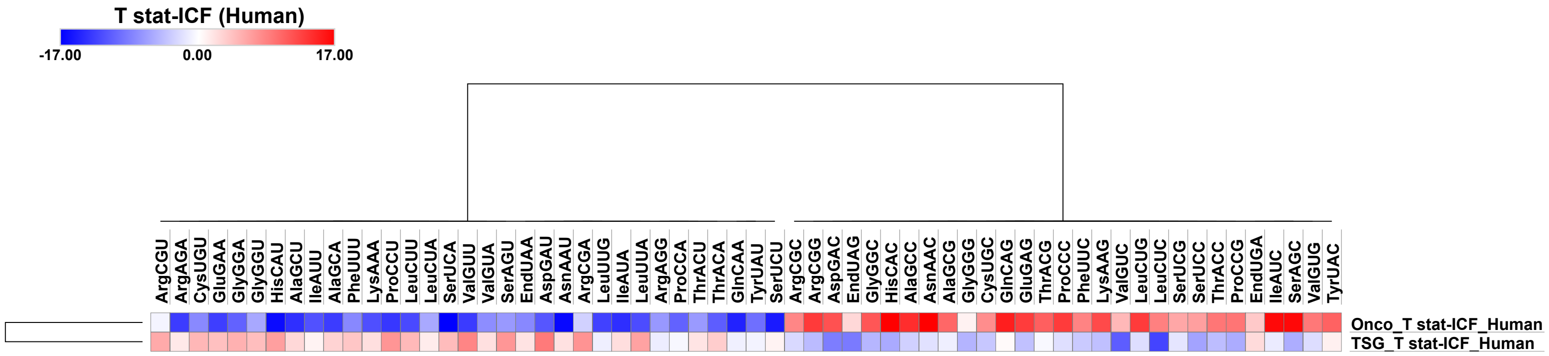

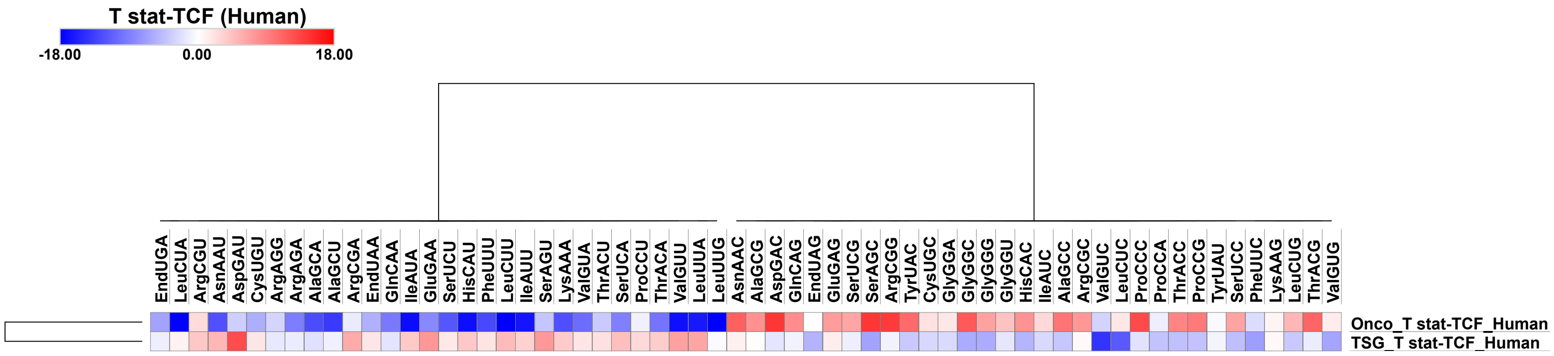

### Supplementary Figure 2

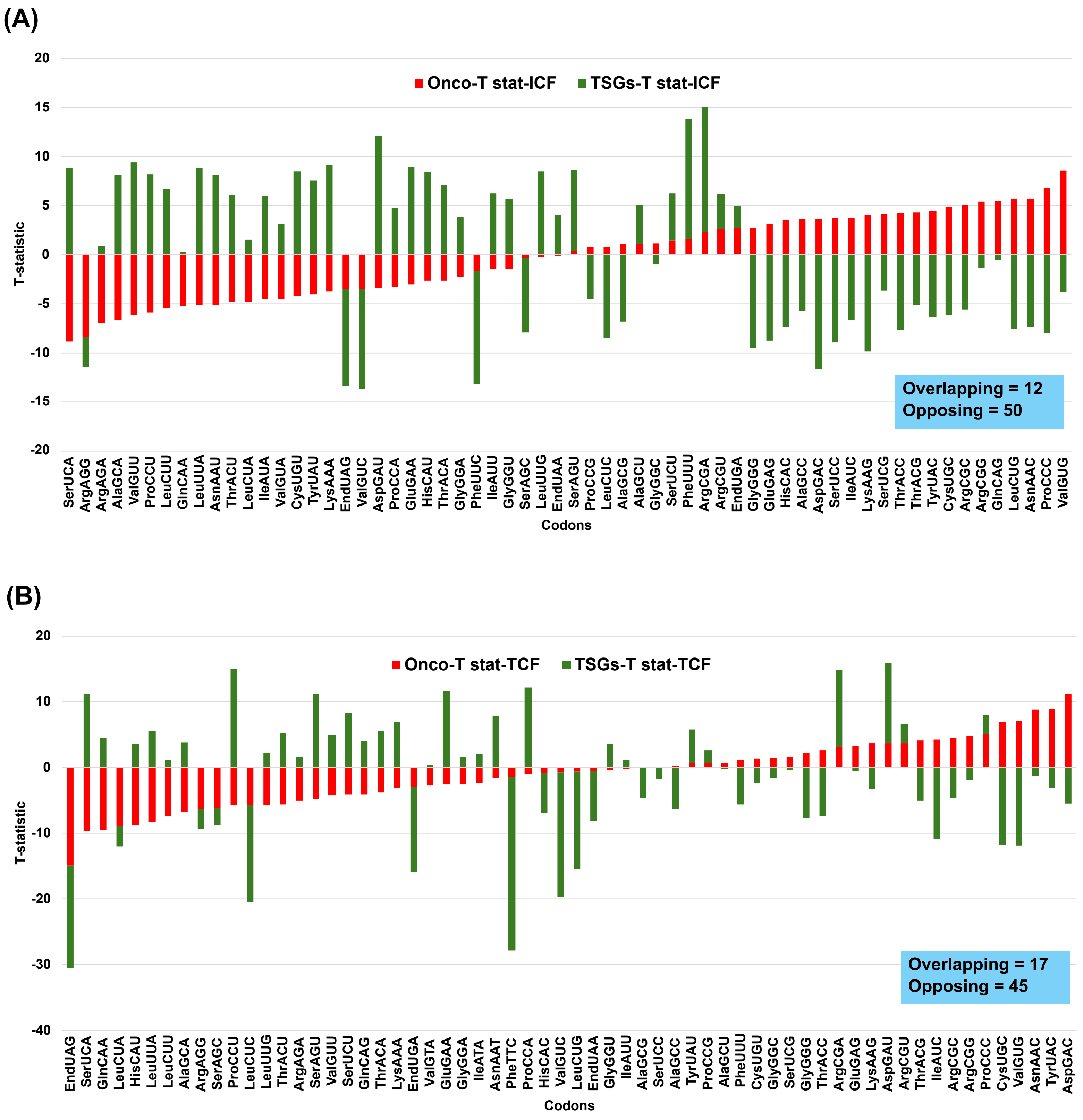

### Supplementary Figure 3

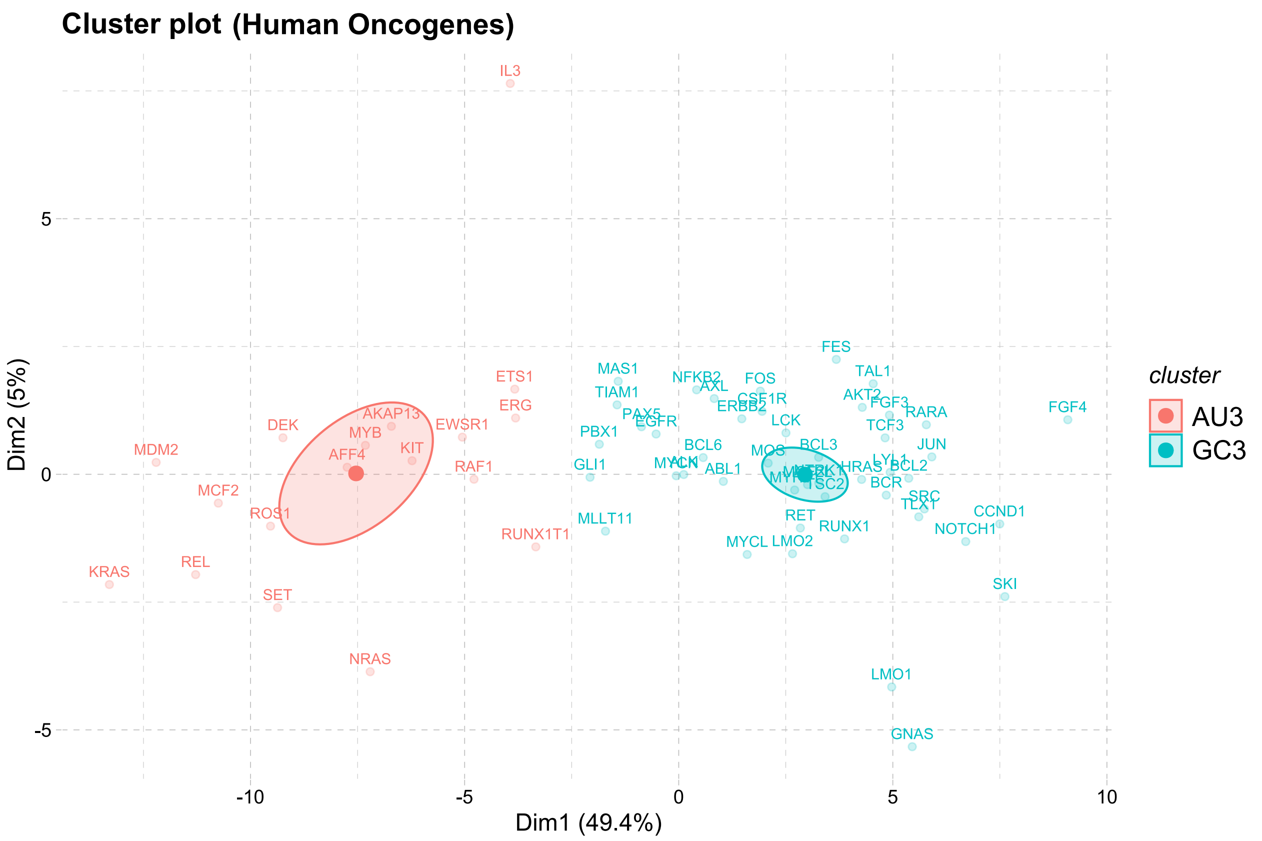

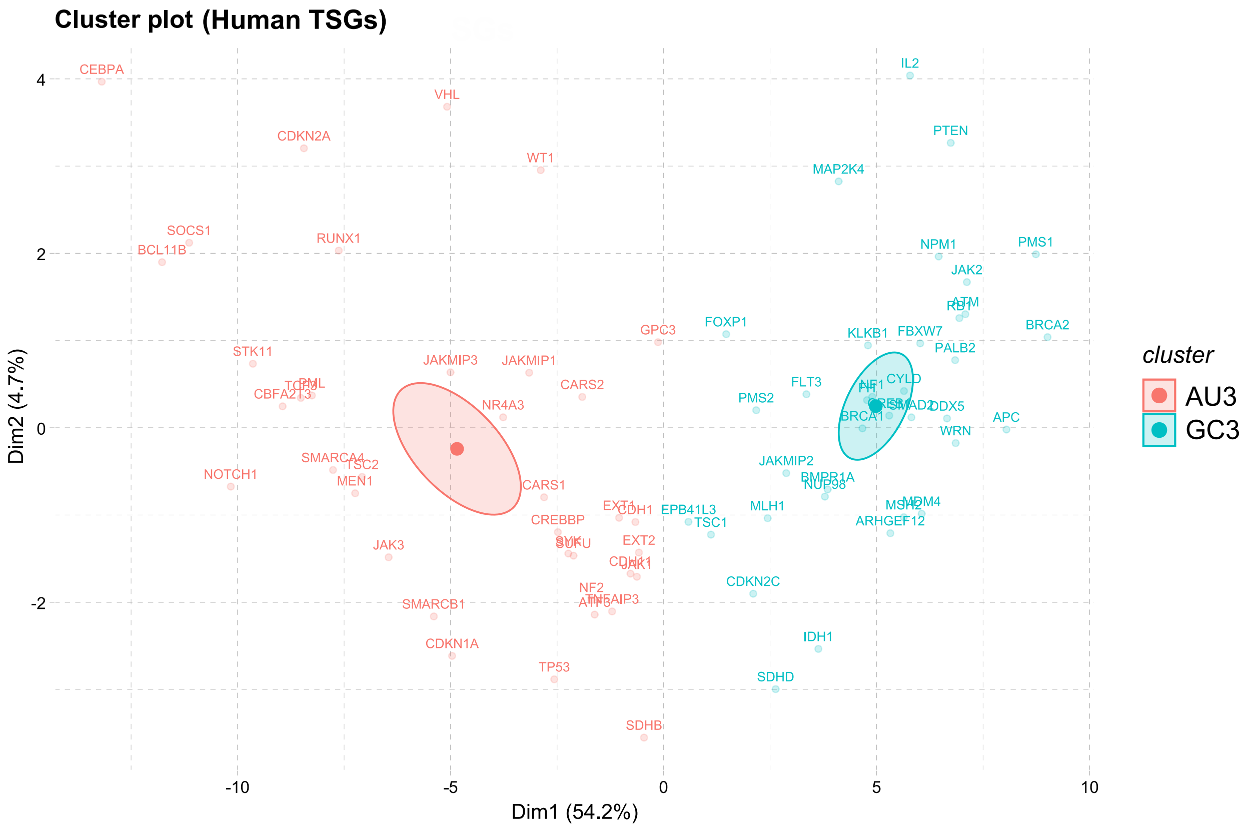

### Supplementary Figure 4

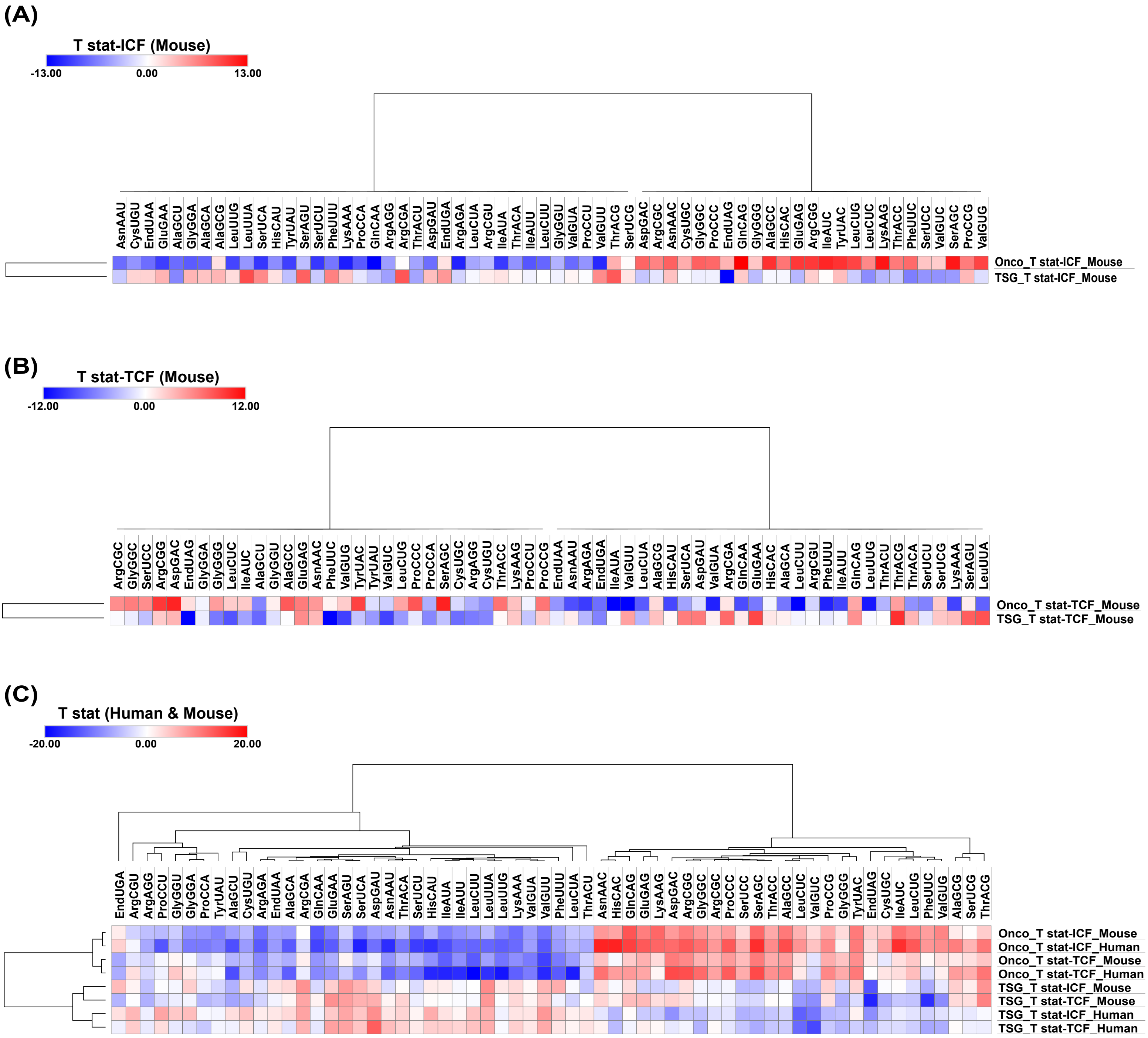

### Supplementary Figure 5

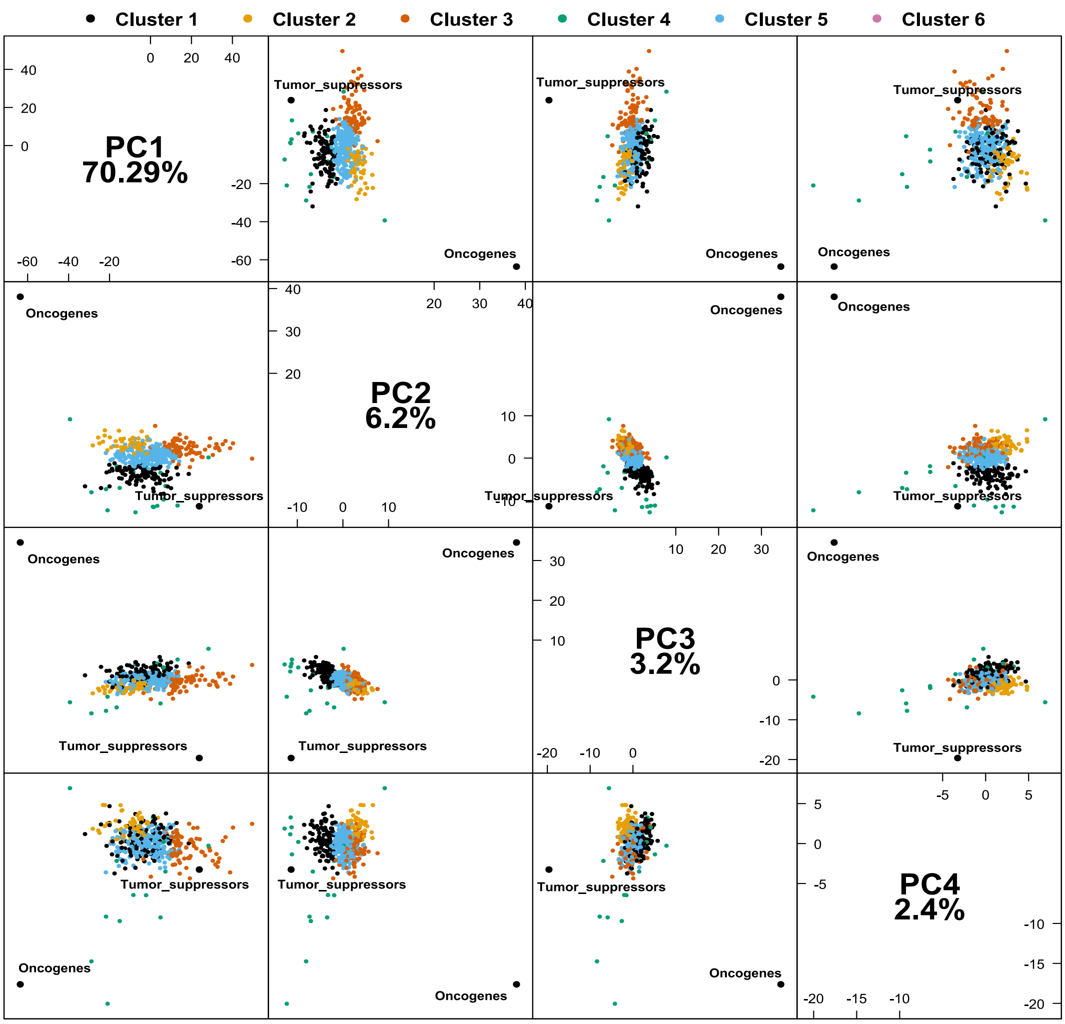

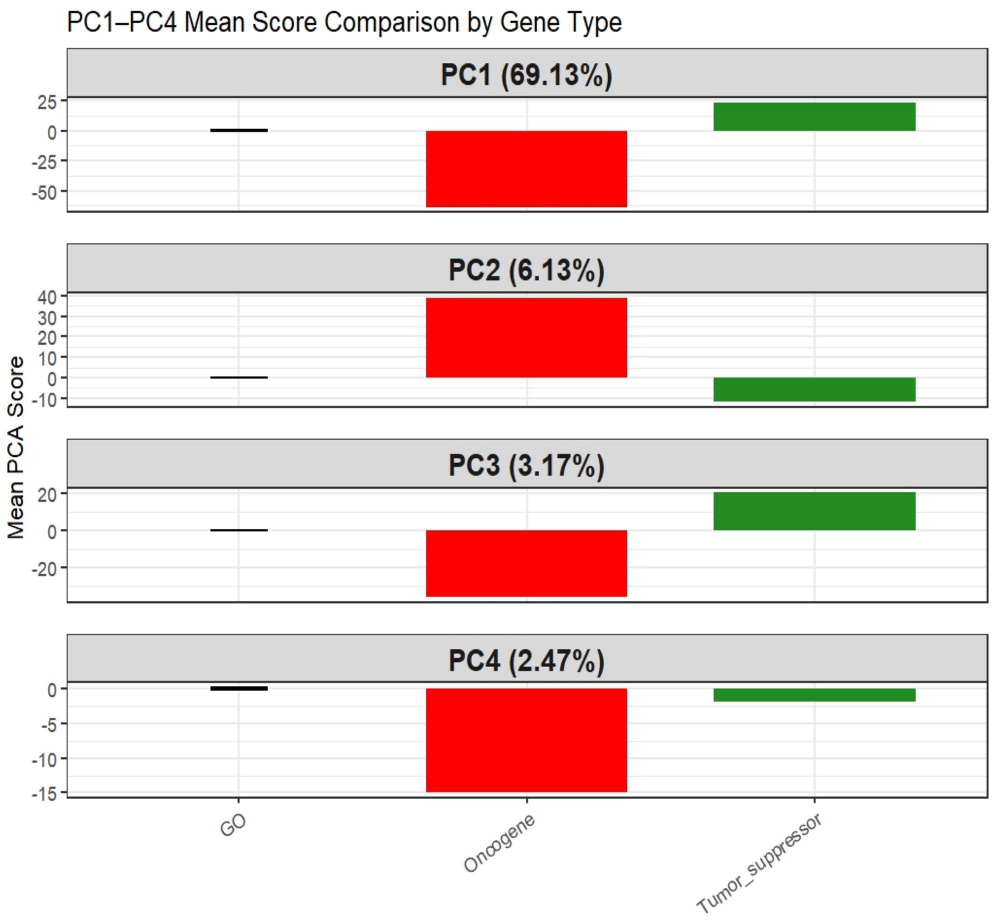

### Supplementary Figure 6

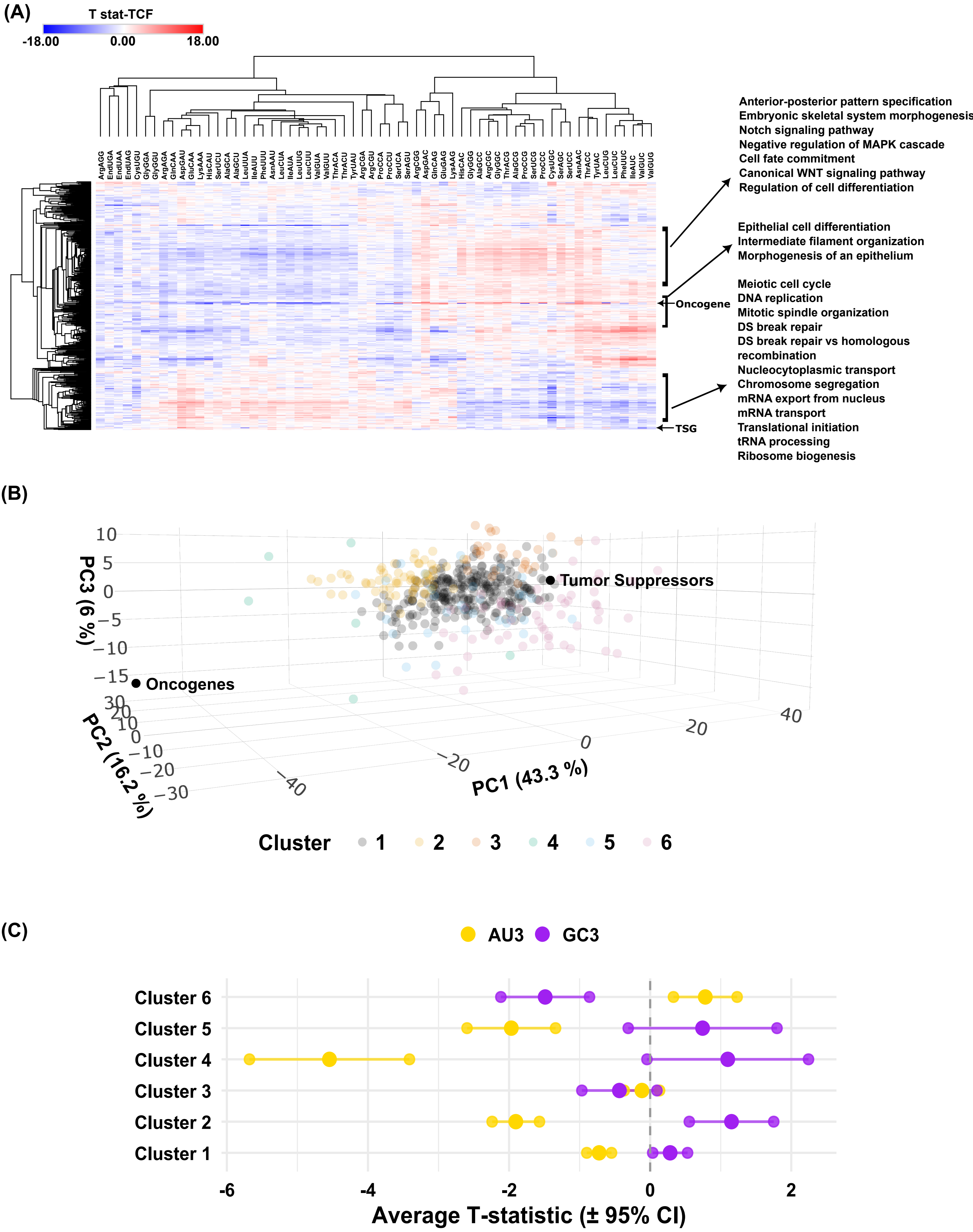

### Supplementary Figure 7

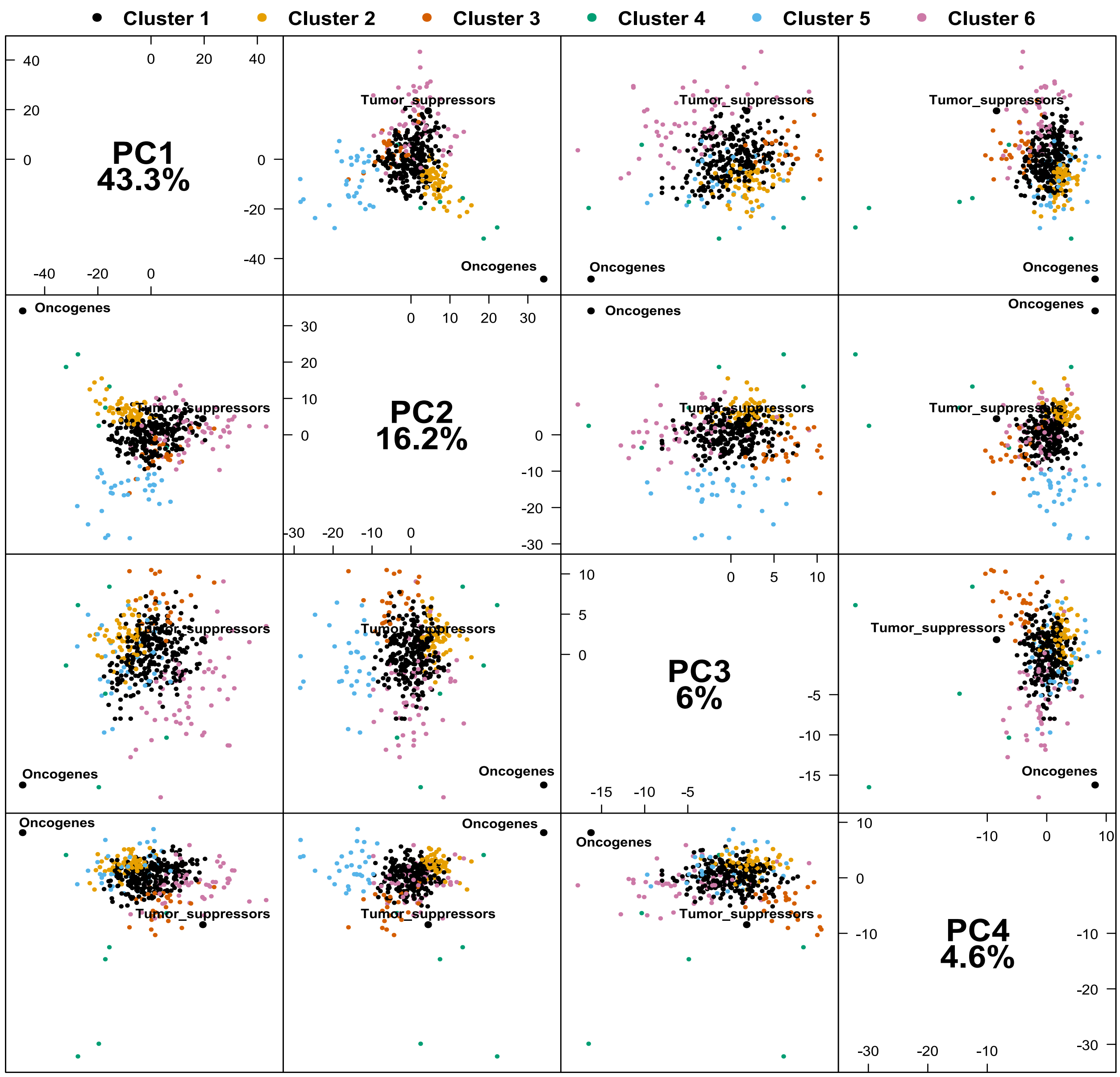

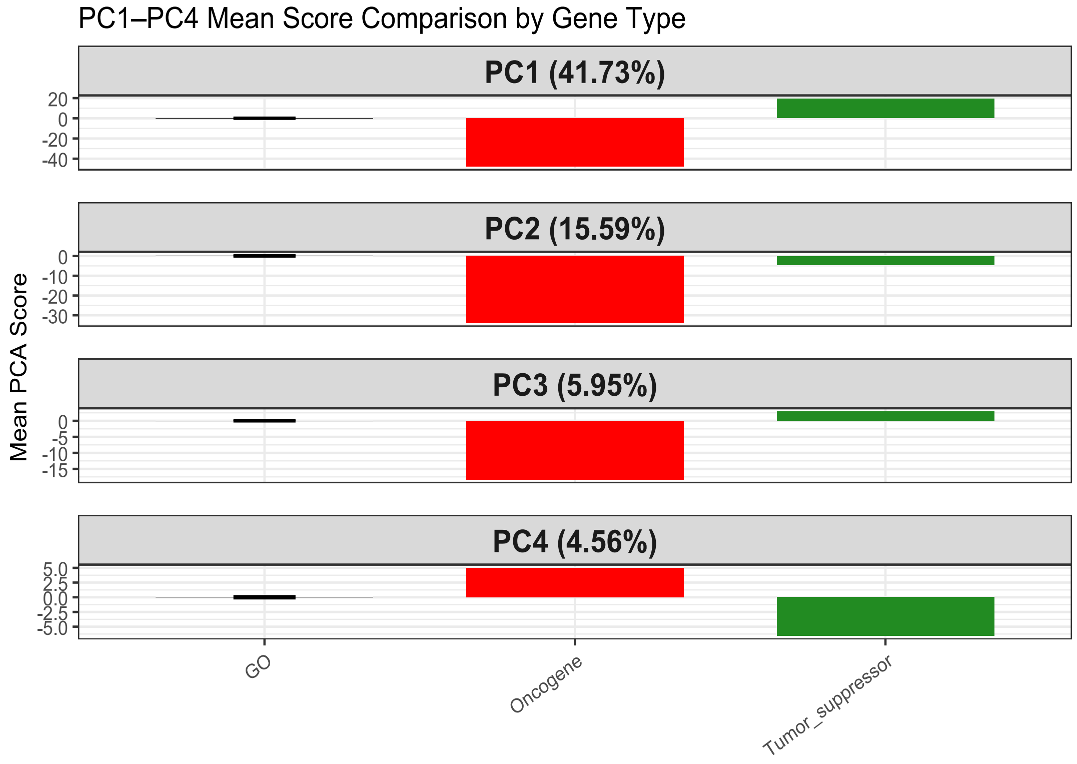

### Supplementary Figure 8

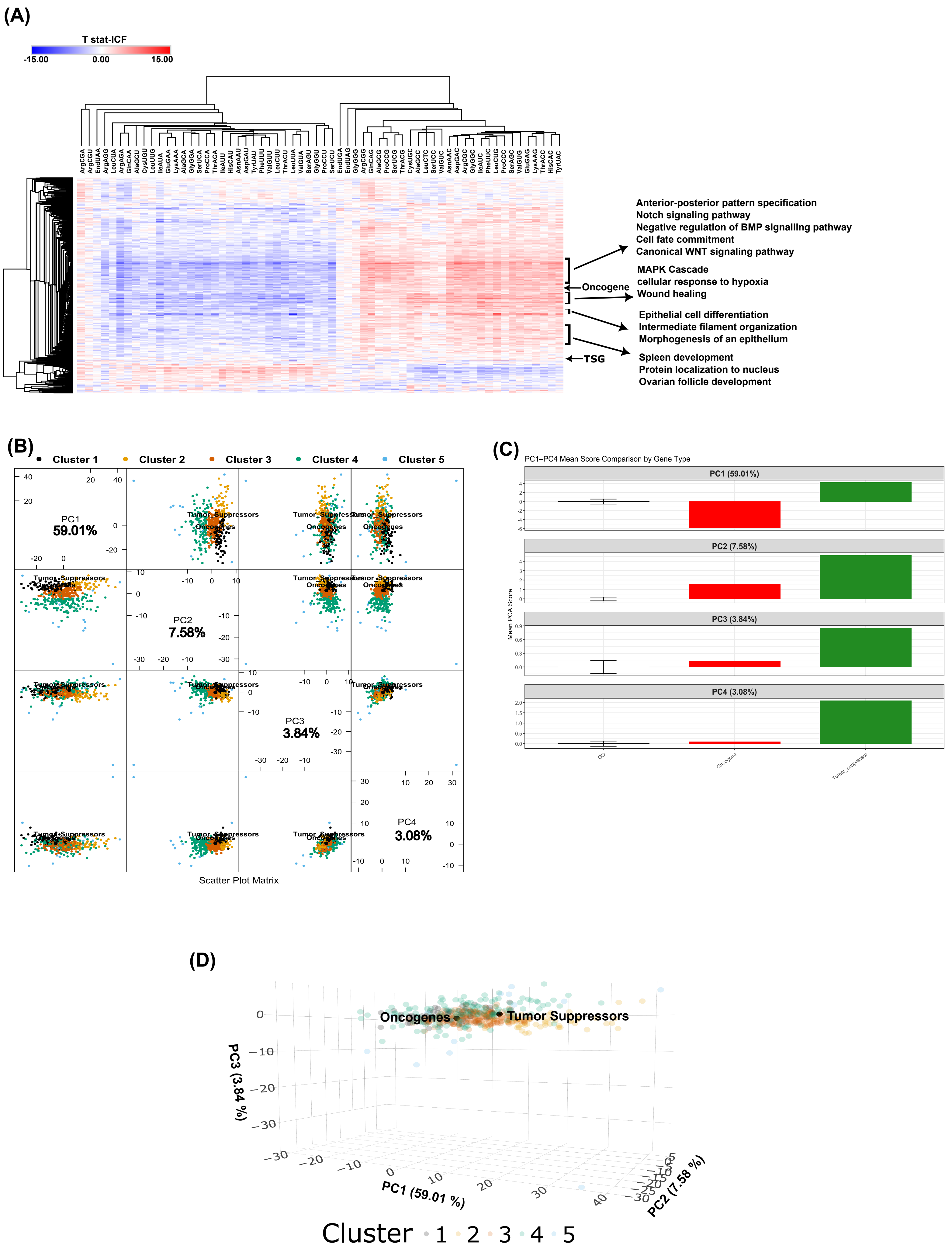

### Supplementary Figure 9

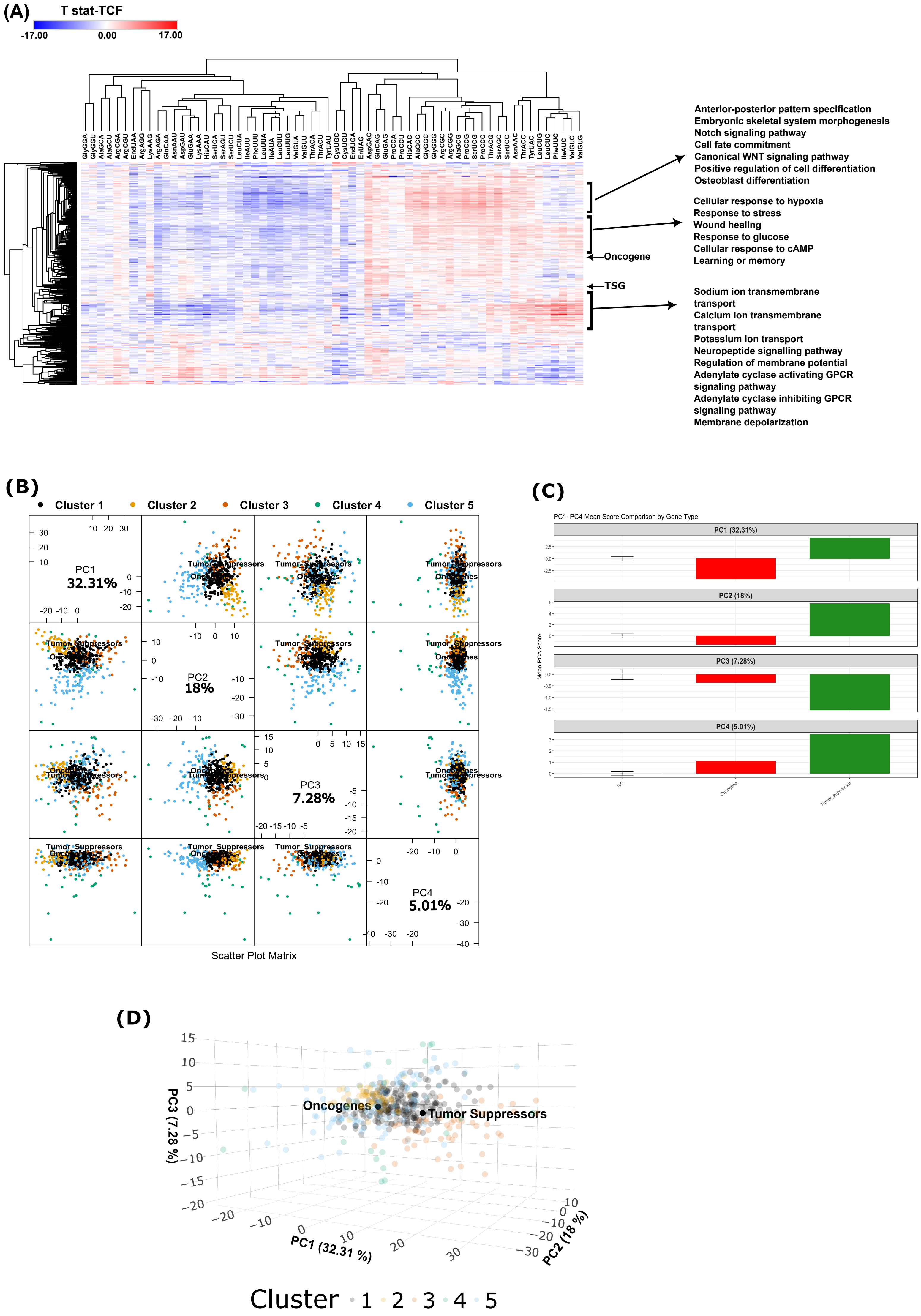
